# The influence of objecthood on the representation of natural images in the visual cortex

**DOI:** 10.1101/2021.09.21.461209

**Authors:** Paolo Papale, Wietske Zuiderbaan, Rob R.M. Teeuwen, Amparo Gilhuis, Matthew W. Self, Pieter R. Roelfsema, Serge O. Dumoulin

**Affiliations:** Department of Vision & Cognition, Netherlands Institute for Neuroscience (KNAW), 1105 BA Amsterdam, Netherlands; Momilab Research Unit, IMT School for Advanced Studies Lucca, 55100 Lucca, Italy; Spinoza Centre for Neuroimaging, 1105 BK Amsterdam, Netherlands; Department of Integrative Neurophysiology, VU University, 1081 HV Amsterdam, Netherlands; Department of Psychiatry, Academic Medical Centre, Postbus 22660, 1100 DD Amsterdam, Netherlands.; Department of Experimental and Applied Psychology, VU University Amsterdam, Amsterdam 1181 BT, Netherlands; Department of Experimental Psychology, Helmholtz Institute, Utrecht University, 3584 CS Utrecht, Netherlands

## Abstract

Neurons in early visual cortex are not only sensitive to the image elements in their receptive field but also to the context determining whether the elements are part of an object or background. We here assessed the effect of objecthood in natural images on neuronal activity in early visual cortex, with fMRI in humans and electrophysiology in monkeys. We report that boundaries and interiors of objects elicit more activity than the background. Boundary effects occur remarkably early, implying that visual cortical neurons are tuned to features characterizing object boundaries in natural images. When a new image is presented the influence of the object interiors on neuronal activity occurs during a late phase of neuronal response and earlier when eye movements shift the image representation, implying that object representations are remapped across eye-movements. Our results reveal how object perception shapes the representation of natural images in early visual cortex.

## Introduction

The visual scenes that we perceive are filled with objects. We readily identify the extent of the objects and their boundaries, a perceptual organization process that is important for our understanding of an image’s meaning. Accordingly, the judgments of people who are asked to mark regions occupied by objects and their boundaries are highly consistent^1^. Object and boundary perception even influence low-level vision, because image elements at object boundaries are better perceived than image elements at less relevant image locations^2, 3^. Furthermore, image elements of objects have a higher perceived contrast than those that are part of the background^4^. Despite these influences on low-level visual perception, it is not yet well understood how objecthood influences neuronal representations in early visual cortex^5^.

Classical descriptions of the activity of neurons at the early levels of the visual system focus on the features that drive neurons, such as the contrast and orientation in a neuron’s receptive field (RF). In addition, there are also non-classical, contextual influences on neuronal activity, which originate from outside the neurons’ RFs and play a role in the grouping of features into objects. Here we focus on two such effects: boundary modulation (BoM) related to the detection of object boundaries, and object-background modulation (OBM) related to the grouping of object features into objects and their segregation from the background.

Neurons in the primary visual cortex (V1) and area V4 increase their firing rate when their RF is centered on an elongated contour that extends well beyond their RF (Fig. 1a)^6, 7^. Elongated contours are relevant for perceptual organization because they usually signal the borders of objects in natural scenes, whereas shorter contours are more likely to be part of the background^8, 9^. BoM is the extra activity elicited by contours that demarcate object boundaries (Fig. 1a,d). Similarly, V1 and V4 neurons exhibit stronger responses when their RF falls on the interior of a perceptual object than when it falls on the background (Fig. 1b)^10–12^. This OBM occurs for all image regions that are part of an object, suggesting that the response enhancement could cause the binding of the distributed representation of features in early visual cortex into coherent perceptual objects^13^. This view is supported by the finding that objects relevant for behavior elicit stronger OBM than objects that are not, implying a relation between OBM and object-based attention that co-selects all features of a relevant object^11^. BoM and OBM are thought to reflect the recurrent interactions within and across visual areas^14^ that determine the spread of enhanced neuronal activity and thereby the perception of spatially extended objects in a scene^13^.

**Figure 1.**
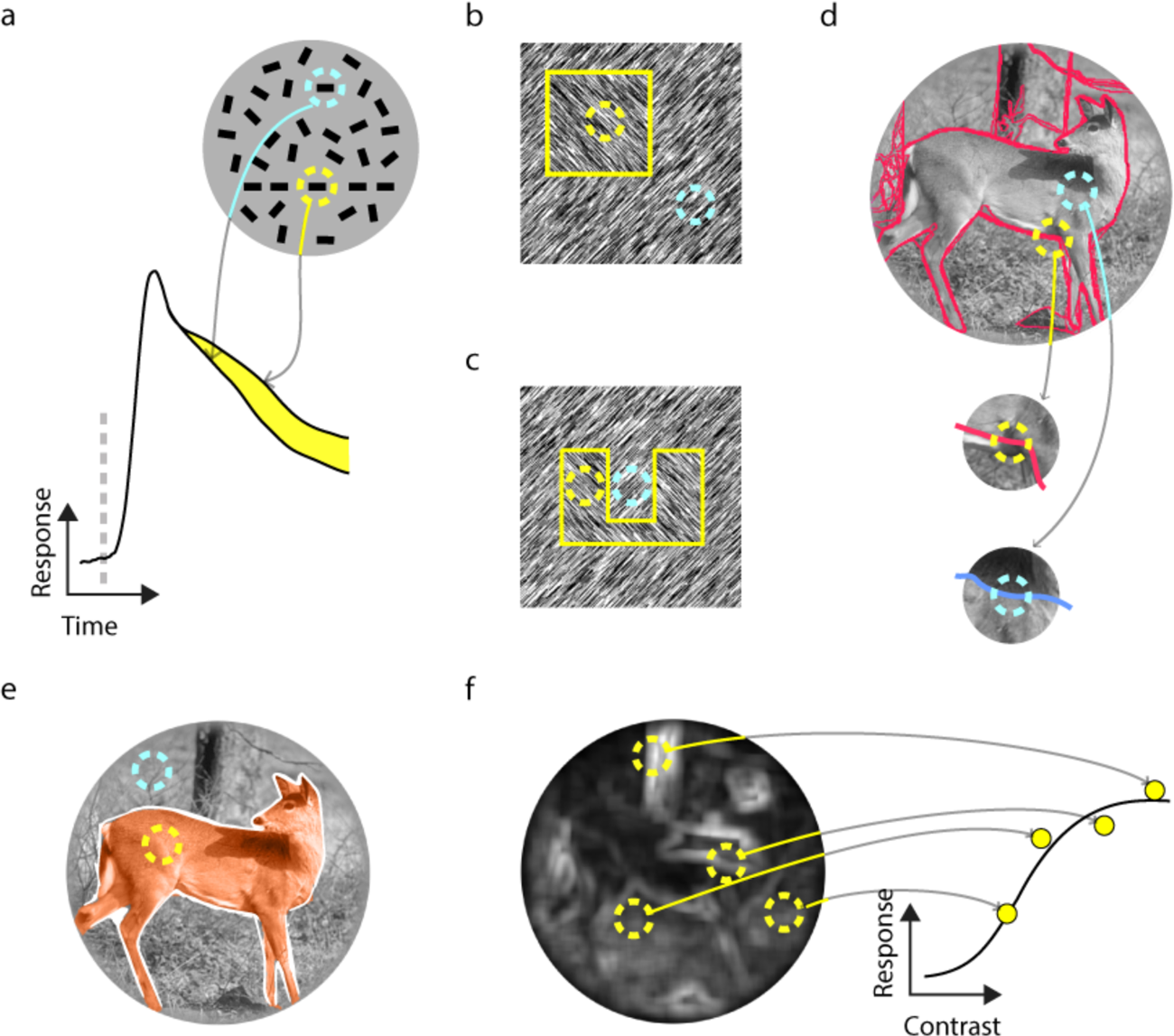
The influence of object perception on neuronal responses in early visual cortex. **a**, The response of V1 and V4 neurons is enhanced when their RF falls on an elongated contour that extends well beyond their RF^6^. **b**,**c,** V1 and V4 neurons exhibit stronger responses when their RF falls on the interior of a perceptual object (square with different orientation in b and ‘u’ shape in c) than when it falls on the background^16^. **d,e**, We ask whether differences between object borders (yellow in d) and other image regions (cyan in d) and between the interiors of objects (yellow in e) and the background (cyan in e) in natural images influence the response of visual cortical neurons. **f**, We compared the response amplitudes evoked by image elements of objects and the background, taking the local contrast in the (p)RF into account.

So far, BoM and OBM have only been measured with artificial stimuli, such as textures and displays with many line elements (Fig. 1a-c). Establishing the relevance of these signals for natural vision is challenging, yet important, because neuronal response properties that do not play a role in the perception of natural stimuli are likely to be of limited relevance^15^. A recent study explored contextual signals in V1 elicited by more complex shapes, such as the texture-defined ‘U’ of Figure 1c^16^, but researchers have, to our knowledge, not yet examined BoM and OBM with natural visual stimuli. If BoM occurs in natural images, we predict that the more salient object boundaries elicit stronger neuronal activity than image elements of the background. Similarly, if OBM occurs for natural images, extra activity should be elicited by object interiors compared to the background.

To investigate the influence of objecthood on neuronal activity in early visual cortex, we used the Berkeley Segmentation data set (BSD), a library of natural images in which human observers marked object boundaries^1^. We used functional MRI to examine neuronal responses across many regions of visual cortex in humans and we also recorded multi-unit activity in V1 and V4 of monkeys to gain insight into the temporal profile of spiking activity. We report that objecthood influences neuronal activity. Object boundaries increased the early neuronal responses and object interiors enhanced activity during a later phase of the response. When subjects made eye movements across the images, these contextual effects carried over from one fixation to the next, implying that objects are remapped across eye movements in early visual cortex^17, 18^.

## Results

### Objecthood modulates responses in human early visual cortex

In the first experiment, we used ultra-high field fMRI at 7 Tesla in four human participants to investigate OBM and BoM within natural images (Fig. 2). Our analysis separated BoM (Fig. 1d) and OBM (Fig. 1e) from the influence of the contrast of image elements by evaluating the influence of image properties in the population receptive field (pRF) on the neuronal responses at each cortical location. We chose 45 images from the BSD (Fig. S1) for which the object boundaries had been annotated by an independent group of human observers^1^.

**Figure 2.**
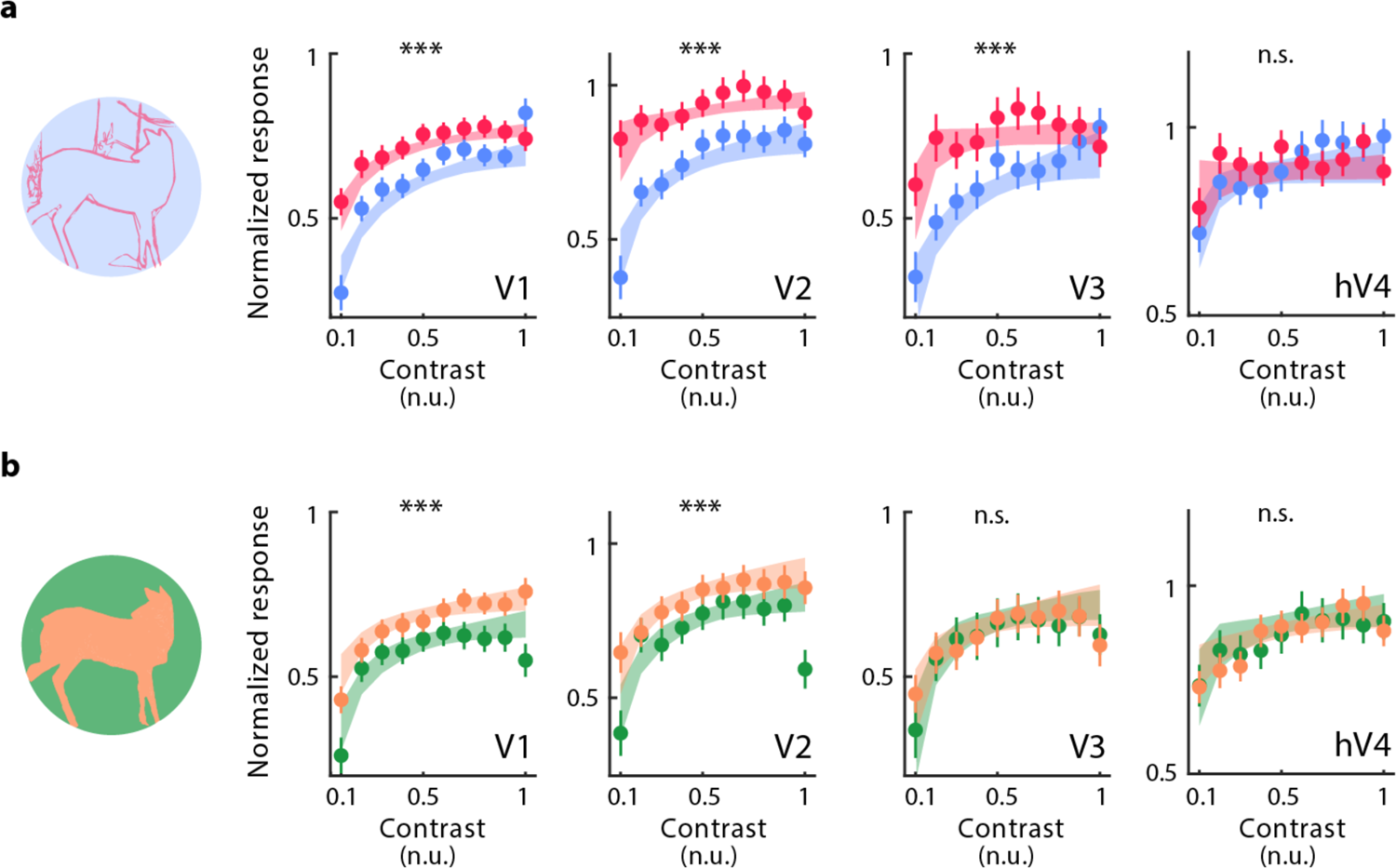
Objecthood influences responses in human early visual cortex. Four participants viewed 45 natural images while their brain responses were recorded with fMRI. **a**, Response amplitudes elicited by object boundaries (red) and background contours (blue) as a function of contrast in the pRF (x axis). **b**, Response amplitudes elicited by object interiors (orange) and the background (green). fMRI responses were normalized to the response to a full-field, 100% contrast stimulus. Shaded regions denote 95% confidence intervals determined by bootstrapping. Bars represent SEM across images (*** indicates p < 0.001, bootstrap test; n.s. non-significant).

To separate the influence of image contrast from object perception we computed the contrast response functions (CRFs; Fig. 1f) for six regions of interest: V1, V2, V3, human V4 (hV4), the lateral occipital visual field maps 1/2 (LO-1/2) and V3-a/b. First, in a separate experiment, we measured the population receptive field for each cortical location (pRF; Fig. S2) ^19^. Second, to compute the CRF, we estimated the response amplitude as function the root mean square (RMS) contrast in each pRF^20^ (10 contrast bins, Fig. 1f).

Next, we computed the CRF separately when the pRF fell on an object border versus a non-border image region, and when it fell an object region versus on the background for every image (Fig. 2b)^1, 21^. We defined object borders in the BSD images as those that were frequently marked by the observers as object boundaries, and contrasted them to non-border image regions that were not marked (Fig S3). These non-border regions could be part of the object interior (as the example in Fig. 1d and Fig. S3) or background (see Methods).

The computation of the CRF allowed us to separate BoM and OBM from the influence of contrast. Cortical responses elicited by object borders were significantly higher than those elicited by non-border image regions in areas V1, V2 and V3 (Fig. 2c; all ps < 0.001, bootstrap test, see Methods), but not in areas V3ab, hV4 and LO-1 and 2 (Fig. S2). Thus, we observed significant BoM in V1-V3. Object boundaries of a particular contrast elicit a larger response, on average, than image regions with the same contrast that do not coincide with object boundaries.

To examine the influence of OBM, we compared CRFs of cortical locations with pRFs on object versus background regions (Fig S3). The response amplitude when a pRF was centered on an object was significantly stronger than when it was centered on the background in V1 and V2 (Fig. 2d; all ps < 0.001, bootstrap test) but not in V3, hV4, LO-1/2 and V3A/b. We observed the same pattern of results also at the level of individual participants, for both BoM and OBM (Figure S4).

We conclude that object borders elicit larger response amplitudes in early visual cortex than non-border image regions, and that object regions elicit more activity than background regions, even if image contrast is the same.

### Objects and their boundaries enhance the spiking activity of V1 and V4 neurons but at different latencies

A limitation of fMRI is its poor temporal resolution and its indirect relation to spiking activity^22^. Therefore, we recorded spiking activity with chronically implanted electrode arrays elicited by BSD stimuli in two macaque monkeys. We placed the arrays in areas V1 and V4 and recorded multi-unit spiking activity (MUA). Whereas the pRFs in the MRI experiment covered the entire images, the RFs of the V1 and V4 neurons in the electrophysiology experiment were confined to a limited region of the visual field. To increase the sample of image patches falling in the RFs, we trained the monkeys to fixate at multiple locations on a total of four images (Fig. 3a).

**Figure 3.**
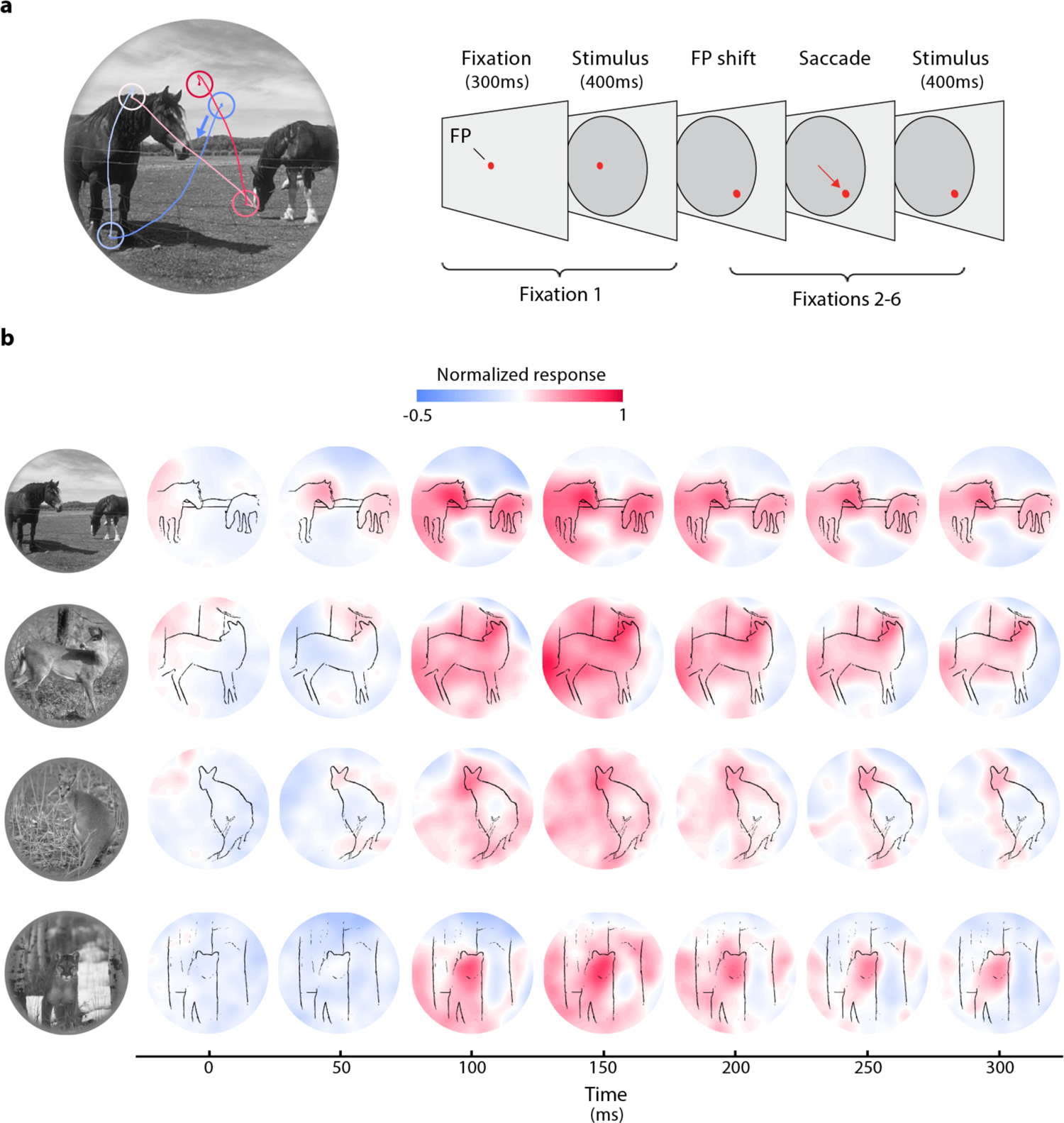
Influence of objects on spiking activity in areas V1 and V4 evoked by natural scenes. **a**, The monkey performed a sequence of eye-movements across the natural images. We presented a natural image once the monkey had maintained gaze on a red fixation point for 300ms. After a delay of 400ms, a new fixation point appeared and the monkey made a saccade to it and maintained fixation for a further 400ms. Per trial, the monkey made a total of 5 eye movements to fixation points sampled from a uniformly spaced grid (∼500 points). **b**, Overlay of V4 spiking activity over the natural images at different time points (average of fixations 2-6). V4 response is determined by contrast, local and global image structure and these factors were disentangled in subsequent analyses. Fig. S3b shows the same analysis for fixation 1 in V4 and fixation 1 and 2-6 in V1.

At the start of the trial, the monkey directed gaze to a fixation point on a gray background. We presented the image once the monkey had maintained fixation for 300ms. After a delay of 400ms, we presented a new fixation point and the monkey made a saccade to it and maintained fixation for a further 400ms (Fig. 3a). The repositioning of gaze was repeated for a total of 6 positions (sampled from a uniformly spaced grid with ∼500 points) per trial. The neuronal response elicited by the image appearing at the first gaze position differs from that for later fixations because the image suddenly appears in the RF. Later fixations are preceded by saccades causing a rapid movement of part of the image through the RF. Furthermore, the image is now familiar and the monkeys may have recognized and segmented the objects during previous fixations. We therefore separately analyzed the response elicited by stimulus onset (Fixation 1) and later fixations (Fixations 2-6). The results for fixations 2-6 were comparable and we therefore pooled the data across these fixations (Fig. S5).

We first examined the overall level of activity in V1 and V4 elicited by the four natural images (Fig. 3b and S3). The response profiles suggest that extra activity is focused on the objects, but the influences of objecthood and contrast were not yet separated in this analysis. To disentangle the influence of BoM and OBM from that of contrast, we determined CRFs by binning the contrast in the RF of neurons in area V1 (77 recording sites, 44 in monkey B and 33 in monkey M) and V4 (22 sites in monkey B), separately for contours that demarcate object boundaries and those that do not. Object borders elicited stronger spiking activity (time-window 0-300ms) than non-object image regions with the same contrast in V1 and V4 (Fig. 4a). BoM occurred during the first fixation as well as during later fixations (all ps < 0.001, bootstrap test) and was present at many V1 recording sites in monkey B (fixation 1, 66% of the sites; fixation 2-6, 77%; Fig. S6) and monkey M (fixation 1, 45%; fixation 2-6, 51%) and at V4 recording sites as well (monkey B, fixation 1, 45%; fixation 2-6, 45%). We determined BoM latency by fitting a curve to the difference in activity elicited by the object borders and non-border image regions, averaged across contrast bins. We estimated latency as the time-point at which the fitted function reached the 33% of its maximum (see Methods)^11, 12, 23^. In V1, the BoM latency was 50ms in both fixation 1 and in later fixations. BoM latency was not significantly different from the latency of the visually driven response (49ms for fixation 1 and 29ms for later fixations), neither for fixation 1 nor for the later fixations (both ps > 0.05, bootstrap test). The same was true for V4 in both conditions (BoM: 59ms for fixation 1; 49ms for fixations 2-6; onset of response: 61ms for fixation 1; 54ms for fixations 2-6; both ps > 0.05, bootstrap test). We cannot directly compare latencies between fixation 1 and later fixations, because in fixation 1 the image replaced a grey background whereas the image moved through the receptive fields preceding the later fixations. We corrected for this difference by computing *Lat_BoM-Vis_*, the difference between the BoM latency and the visual latency and compared it between fixation 1 and later fixations across recording sites (Fig. S7). *Lat_BoM-Vis_* did not differ between fixation 1 and fixations 2-6 (p > 0.05, Wilcoxon signed-rank test; Fig. S7).

**Figure 4.**
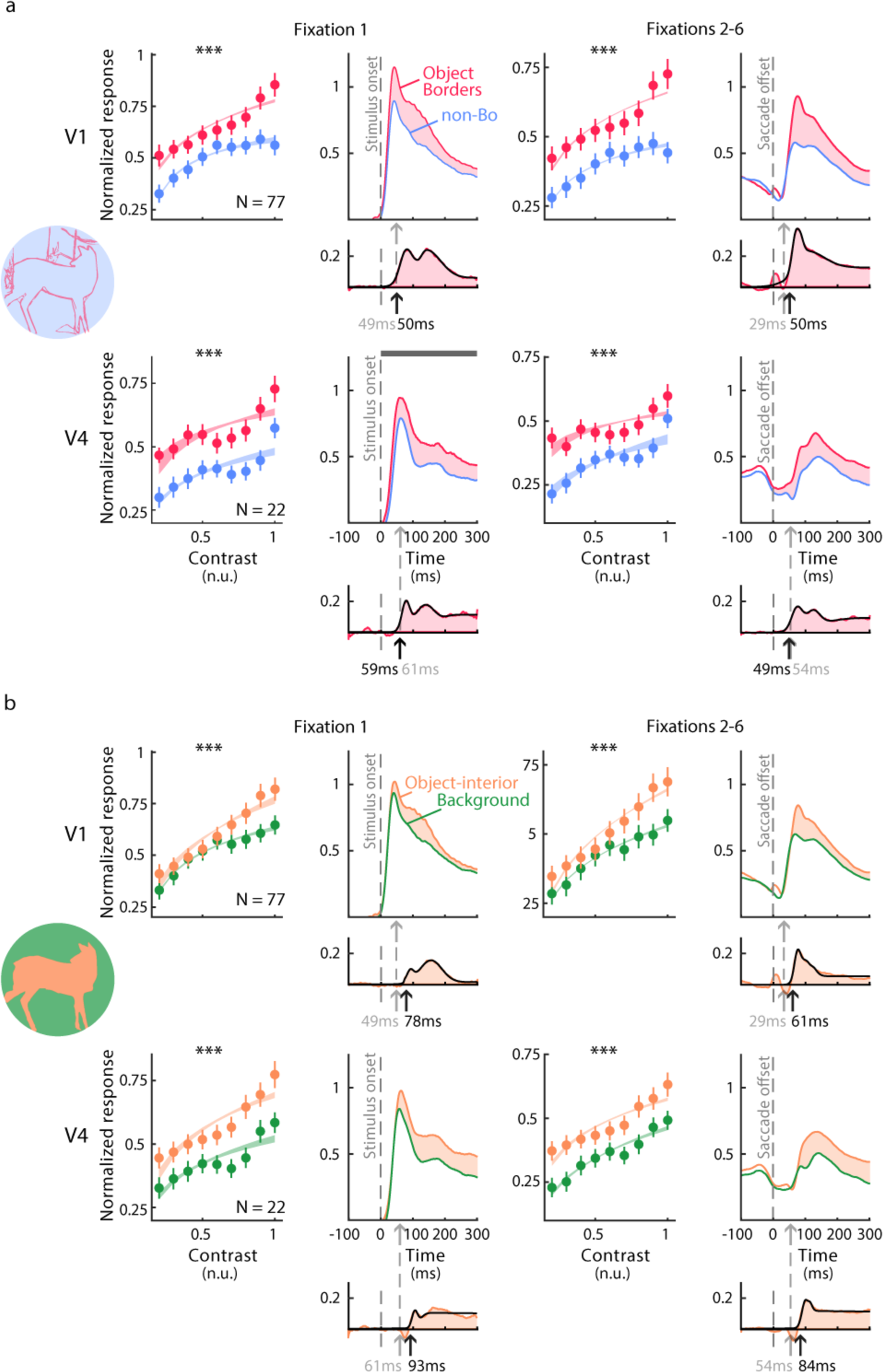
Object borders and their interiors enhance the spiking activity of V1 and V4 neurons relative to background regions. **a**, Average CRFs (left) and MUA time-courses (averaged across contrast bins; right) in V1 (top row) and V4 (bottom row) for object-borders (red) and non-border image regions (blue). BoM is significant in both areas. Shaded regions around the CRFs denote 95% confidence intervals (determined by bootstrapping). Error bars indicate SEM across recording sites (***, p < 0.001, bootstrap test). **b**, CRFs and MUA time-course elicited by the object-interior (orange) and background (green). Black arrows indicate the latency of either BoM/OBM and gray arrows the latency of the visually driven response.

Next, we compared the response elicited by object interiors to that elicited by background regions (Fig. 4b). Regions that were part of objects elicited more activity in V1 and V4 than background regions, both during the first fixation and later fixations (all ps < 0.001, bootstrap test). Many V1 recording sites exhibited OBM (monkey B, fixation 1, 41%, fixations 2-6, 52%; monkey M, fixation 1, 67%, fixations 2-6, 36% of sites with p<0.05, bootstrap test) and OBM was also present in V4 (monkey B, fixation 1, 41% of sites in V4, fixations 2-6, 45%). Hence, OBM also occurs for natural images: image elements of objects elicit a stronger activity than those that are part of the background.

In V1, the latency of OBM during the first fixation was 78ms, which was later than the onset of the visually driven response (p < 0.05, bootstrap test). We next examined whether the OBM latency was shorter for later fixations, because the monkeys may have segmented the image in figure and background during the previous fixations. Interestingly, the median OBM latency across sites for fixations 2-6 was only 61ms, and not significantly different from the visually driven response and BoM (p > 0.05, bootstrap test). To correct for the difference in visual stimulation we computed *Lat_OBM-Vis_*, the difference between the latency of OBM and the visual response across recording sites. In V1, *Lat_OBM-Vis_* was 13ms shorter during fixations 2-6 than during fixation 1 (p < 0.05, Wilcoxon signed-rank test; Fig. S7). In V4, the latency of OBM was later than the onset of visually driven response both for fixation 1 and fixations 2-6 (both ps < 0.05, bootstrap test). The difference in *Lat_OBM-Vis_* between the fixation periods was not significant, but we cannot exclude the possibility that this was caused by the smaller number of V4 recording sites.

The earlier OBM in V1 during fixations 2-6 suggests that it may have carried over from earlier fixations during which the monkeys had already recognized and segmented the objects, in accordance with previous studies demonstrating that image segmentation results can be remapped across saccades^17, 18^.

BoM entails a comparison of the response elicited by object borders and other image elements, which were inside the objects or in the background, whereas OBM entails a comparison of image elements inside objects to background elements (Fig S3). BoM and OBM are not independent because both measures compare neuronal activity evoked by some of the object elements and background elements. We therefore also investigated the amount of unique variance in the activity of V1 and V4 neurons (time window 0-300ms) explained by BoM, OBM and contrast (Fig. 5a; see Methods). Each predictor explained a significant amount of unique variance in both areas and both fixation conditions (all ps < 0.001, t-test). In V1, contrast explained 41.0% of the variance, BoM 8.3% and OBM 5.5% during the first fixation and the values increased slightly to 44.8%, 10.1% and 6.4% for fixations 2-6, respectively. In V4, contrast explained 19.0%, BoM 21.4% and OBM 16.0% of the variance during the first fixation and these values were 20.8%, 21.1% and 35.0% for the later fixations. Hence, BoM and OBM accounted for a significant fraction of the explainable variance. Contrast explained less variance in V4 than in V1 whereas the contributions of BoM and OBM were larger in V4. It is of interest that the fraction of variance explained by OBM in V4 increased from 16% for the first fixation to 35% for the later fixations. This result suggests that the extra activity elicited by the interior of objects builds up across between the first presentation of the image and subsequent eye movements, possibly because the scene is already known.

**Figure 5.**
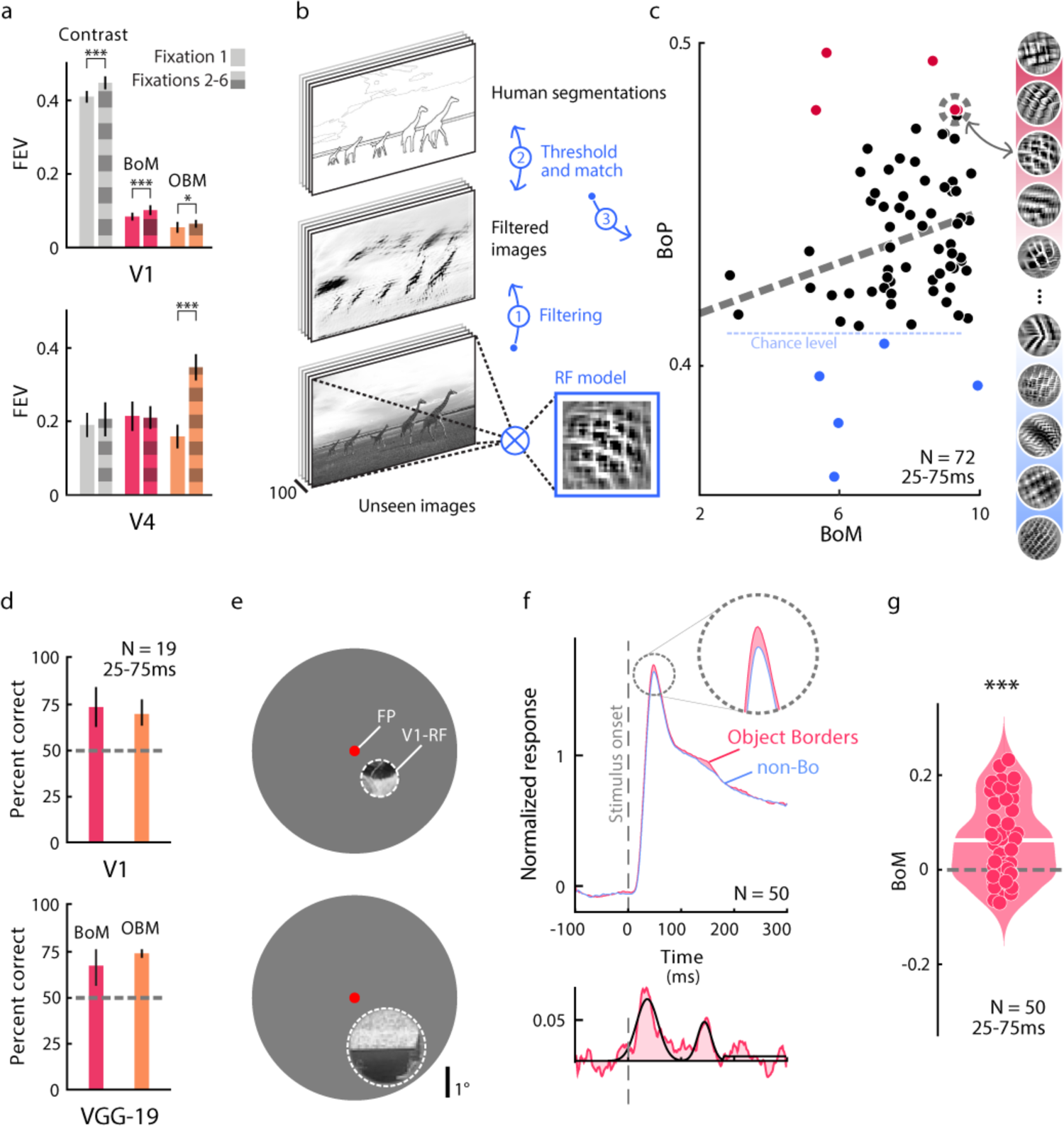
Explained variance in V1 and V4 and V1 tuning during the onset response. **a**, Fraction of variance of V1 and V4 activity explained by contrast, BoM and OBM (0-300ms time window) for fixation 1 and fixations 2-6 (***: p < 0.001; *: p < 0.05, Wilcoxon signed rank). Error bars denote SEM. **b**, We derived RF models for each V1 recording site using ANNs, and applied them to a separate annotated set of natural images to examine how well they can detect object borders. We calculated a measure of border-detection performance (BoP) for every V1 recording site. Step 1, applying the V1 tuning to the image. Step 2, thresholding of activity and correlation with human judgements. Step 3, measurement of BoP of the V1 recording site. **c**, Correlation between BoM (x-axis) and BoP (y-axis) across V1 recording sites (p < 0.05, parametric test). Blue dashed line, significance threshold for border detection (p < 0.05, permutation test). **d**, Accuracy of binary classifiers of object contours (red bar) and interiors (orange bar) based on the early response (25-75 ms) of 19 V1 sites (upper panel) or the VGG-19 conv3_1 layer (lower panel). Classifiers detected object contours and interiors above chance level. Error bars denote 95% confidence intervals (determined by bootstrapping). **e**, Example isolated BSD image patches matching RFs of different V1 recording sites. **f**, Time-course of the V1 responses. Object borders elicited stronger early activity than non-border image patches. **g**, Distribution of early (25-75ms) BoM elicited by image patches across recording sites (white bar indicate the median BoM; ***, p < 0.001, Wilcoxon signed-rank test).

### The tuning of the early V1 response is selective for object borders

We were surprised to find BoM in V1 at a latency of 50ms (Fig. 4a), because it is much earlier than the latency of ∼95ms typically observed for elongated contours in synthetic images^7^. The longer BoM latency of previous studies is compatible with an effect of feedback from higher cortical areas to V1, but a latency of 50ms might be too short for such a feedback loop. An important difference between the present approach and previous studies with synthetic images is that we did not equate the features of contours that form the boundaries of objects and those that were in the background, even though contrasts were matched. We therefore hypothesized that object contours in natural images have other features, on average, than background contours^9, 24^, which could explain the early BoM. In other words, some V1 neurons might be tuned to the features of object contours and extract them in their feedforward response, driven from within the RF.

We exploited recent advances in artificial neural networks (ANNs) to study the tuning of V1 recording sites and to examine if it can account for the extra activity elicited by object boundaries^25–27^. As a model for V1 tuning we chose layer conv3_1 of VGG-19 (and several other models, Fig. S8), which is the state of the art in predicting V1 responses to natural images^28, 29^, and used a two-stage convolutional mapping to take both the spatial and feature selectivity of neurons at individual recording sites into account (see Methods)^26^. We confirmed previous studies^28, 29^ demonstrating that the ANN approach for the modeling of V1 tuning outperforms previous models (Fig. S8). To gain insight into the tuning of the V1 neurons and visualize their preferred features, according to the ANN model, we examined the synthetic images that maximized the model response. Figure 5c illustrates a few of these synthetic images (for illustration purposes; we did not present these images as stimuli). We then applied these RF models to an independent set of 100 natural images that had been annotated by human observers to examine if they indeed predicted extra activity for object boundaries (Fig. 5b,c and S9). Specifically, we filtered the unseen images with the RF models (step 1 in Fig. 5b) and compared the filtered images to the human annotations (step 2 in Fig. 5b). For every recording site we determined their border detection performance (BoP), a measure that quantifies how well the V1 RF models predict the human-annotated borders (step 3 in Fig. 5b and Y-axis in Fig. 5c). BoP is an accuracy score that takes the uneven class distribution of salient and non-salient borders into account (F-measure, see Methods). Interestingly, 93% of the VGG-19 models of V1 tuning detected objected contours above chance level (ps < 0.05, permutation test), which indicates that V1 tuning is indeed useful for boundary detection.

What is the relation between BoM elicited in V1 by the four pictures of our electrophysiological experiments and the BoP of the same recording sites for a different set of images? We computed the correlation between BoM in the early time-window (25-75ms: x-axis in Fig. 5c) and BoP (y-axis in Fig. 5c), across recording sites. The correlation coefficient was 0.25 (p = 0.037, t-test), indicating that V1 neurons that express BoM at an early latency are tuned to low-level feature differences that discriminate between object and non-object contours (Fig. 5c).

We next examined how much information about object contours is present across the recorded population of V1 neurons. We built a binary classifier based on the early (25-75ms) V1 activity in response to trials with object-contours or elements with the same contrast in the neurons’ RF. To ensure that complex patterns signaling the presence of an object (e.g., the entire head of an animal) could not be detected by the RFs, we only included the 19 recording sites with smallest RFs (<1.5°) in this analysis. Classification of object boundaries during single fixations had an accuracy of 73.5%, which is well above the chance level of 50% (Fig. 5d, top, red bar; p<0.001, bootstrap test). Interestingly, when we used the activity of the entire conv3_1 layer of the VVG-19 ANN to detect object-borders the accuracy was similar (Fig. 5d, bottom, red bar; 66.4%, p<0.001, bootstrap test).

Hence, object-contours can be detected with a reasonable accuracy based on local information in individual RFs. We carried out an extra experiment to confirm that the early BoM reflects V1 tuning rather than a contextual influence. We removed the context by copying circular image patches from the BSD that matched the V1 MUA RFs in size onto a grey background. We chose patches with a similar RMS contrast that did or did not contain an object border and centered them on the RFs of neurons at 50 recording sites in monkey B (Fig. 5e). As predicted, patches with object borders elicited a slightly stronger V1 response than patches without object borders with the same contrast (p < 0.001, Wilcoxon signed rank test; Fig. 5f,g). Hence, the tuning of V1 neurons indeed explains a fraction of the extra activity elicited by object boundaries.

Our finding that BoM is partially explained by V1 tuning begs the question of a possible contribution of V1 tuning to OBM, i.e. the extra activity by the object interior. We therefore also examined low-level differences between image elements of objects and backgrounds and built a binary (linear) classifier to discriminate between object and background regions, based on the early (25-75ms) response of the same 19 V1 recording sites as above, using trials with the same contrast. The classification accuracy during single fixations was 69.9% (Fig. 5d, top, orange bar; p<0.001, bootstrap test) and it was in the same range for the conv3_1 layer of VVG-19 (Fig. 5d, bottom, orange bar; 74.0%, p<0.001, bootstrap test). Thus, even though OBM emerges later than BoM, the activity of a small number of V1 neurons is enough to differentiate between features that characterize the interior of objects and the background.

### Contextual BoM in natural images

The early BoM in natural images is driven by the information in the RF. It differs from BoM in previous studies^6, 7^, in which it was a contextual effect driven by information outside the neurons’ RF. Does BoM also occur for natural images if the image elements in the RFs are kept the same? In a further experiment, we placed the RF of 98 V1 recording sites (68 in monkey B and 30 in monkey M) on object borders and other locations in 12 natural images from the BSD, while keeping the image patch in the RF constant (Fig. 6a). Specifically, we copied an image patch with an object border and pasted it at a background location to create a condition in which the same image patch is not perceived as object border. An example image is shown in Fig. 6a (left panel) where we copied a part of the back of the elephant into the background. On average, the object contours elicited a stronger V1 response than the same image patches presented at background locations (p < 0.001, Wilcoxon signed-rank test across recording sites; Fig. 6b). The latency of BoM in this experiment was 81ms, i.e. it now occurred during the delayed phase of the V1 response.

**Figure 6.**
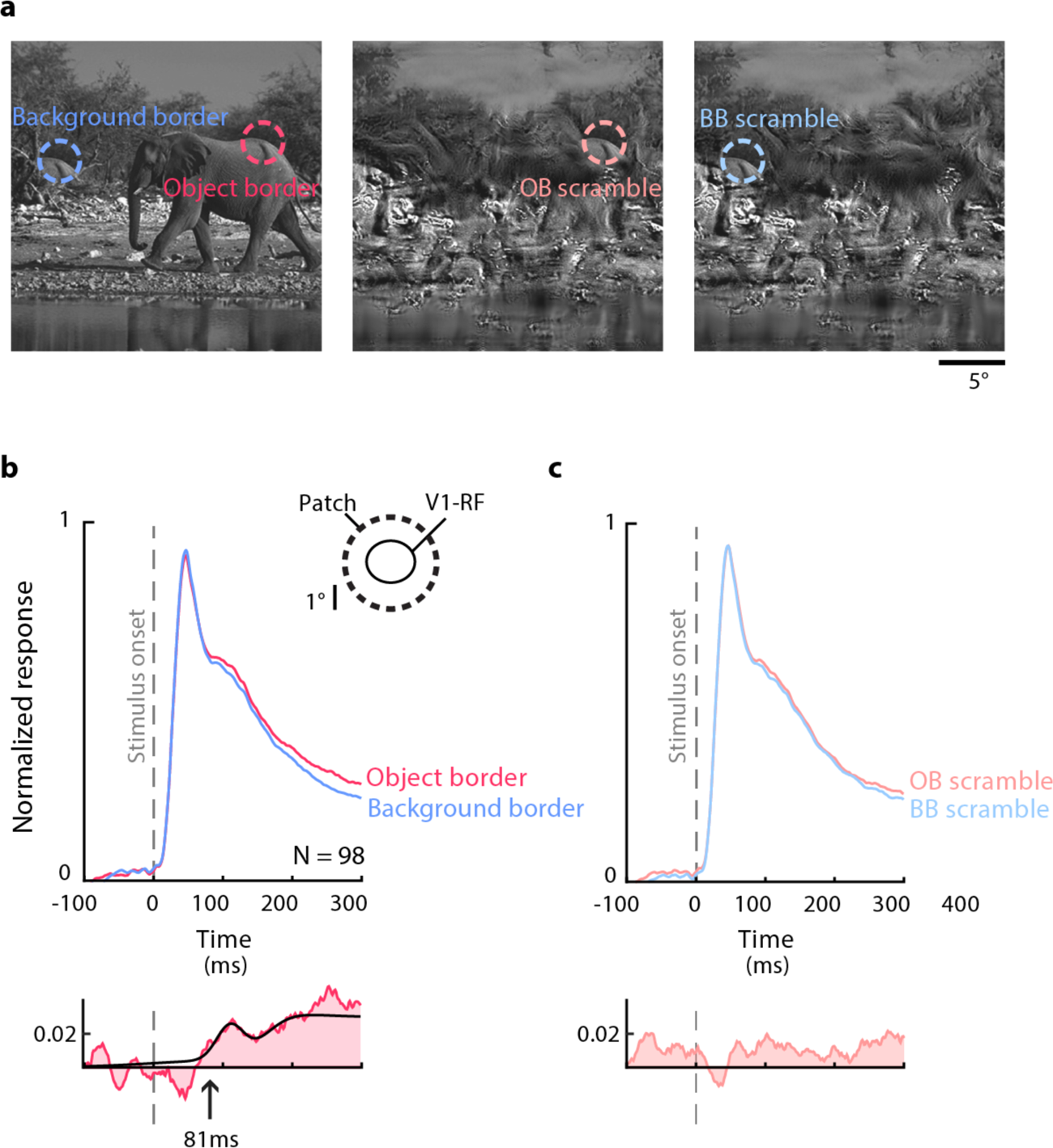
Contextual BoM in V1. **a**, To examine the role of contextual information around the V1 RF, we modified natural images ensuring that the same features were present the RFs. We either copied an image patch with an object border to a background location (left panel) or removed the object from the scene by creating *metamers* (middle and right panel). The RF stimulus was kept constant across all the conditions. **b**, Average V1 response elicited by image regions that demarcated object boundaries (red) or were part of the background (blue). Lower panel, response difference. BoM in this condition had a latency of 81ms. **c**, The responses elicited by the metamers revealed no significant differences.

As a control, we placed the image patch at identical locations of synthetic metamers of these images. The metamers had the same orientations, phases, spatial frequencies, auto- and cross-correlations and marginal statistics, but the layout of objects was scrambled^30^. In the example metamer of Fig. 6a (middle and right panels), the transitions between water, trees and air were at the same locations but the elephant was removed (other example metamers are shown in Fig. S10).

BoM was absent for the metamers (p > 0.05, Wilcoxon signed-rank test). To investigate if the level of BoM differed between the metamers and the original images, we performed a repeated-measures two-way ANOVA with object-border and scrambling (2 levels each) as factors. The main effects of object-borders and scrambling were both significant (salience, F_1,97_ = 28.6, p < 0.001; scrambling, F_1,97_ = 5.42, p = 0.022). Importantly, the interaction was also significant at the population level (F_1,97_ = 6.74, p = 0.011) and at many of the individual recording sites (at p < 0.05; 40% of the sites in monkey B and 73% in monkey M). Hence, if RF stimulus is kept constant, contextual information enhances the V1 activity elicited by object borders, at a latency of ∼80ms. These results, taken together, indicate that there are two processes that jointly explain the enhanced activity elicited by object boundaries. The tuning of V1 neurons enhances their representation from an early time point onwards and the scene context causes an additional activity increase at a longer latency.

## Discussion

We investigated how objects in natural images influence neuronal activity in early visual cortex and observed widespread influences of objecthood on neuronal activity in the human early visual cortex. These results were mirrored by the early and late modulation of neuronal activity in areas V1 and V4 of monkeys. Early influences were related to the tuning of the neurons, causing object boundaries to elicit more activity than background elements. However, if we held the image elements in the RF constant, image elements that were part of an object also elicited more activity than elements that were part of the background. This contextual influence manifested during a later phase of the neuronal response, which suggests the involvement of feedback from higher areas and/or horizontal interactions within visual areas. Whereas previous studies on figure-ground segregation and contour integration in early visual cortex used well-controlled, but artificial stimuli, the present results demonstrate that these findings generalize to natural vision. The results are in accordance with theories proposing that image elements of figures are labeled by enhanced neuronal activity in early visual cortex to segregate them from the background^10, 13, 31^.

Despite the different tasks and recording modalities between humans and monkeys, the neuronal responses in V1 were strikingly similar between the two species (Fig. S2). Our fMRI experiment revealed that BoM for natural images is present in V1 and other areas of early visual cortex. Object regions evoked stronger response than backgrounds in areas V1 and V2 but OBM did not reach significance in a number of higher areas, including hV4. In contrast, in the electrophysiological experiments in monkeys, OBM was present in V1 but even stronger in V4. This discrepancy may be related to differences between species, experimental setups and differences in spiking versus fMRI measures of neural activity^22, 32^. Another relevant difference is the larger size of fMRI pRFs in hV4 compared to neurophysiologically determined V4 RFs. The larger pRF sizes in hV4 may include more neurons with RFs not on the boundary and thereby dilute the BoM signal. Human fMRI allowed us to link the neural responses to human perception, and the monkey neurophysiological experiments allowed us to measure the timing of BoM and OBM and relate it to previous neurophysiological work with synthetic stimuli.

### Early and later object boundary signals

Unexpectedly, natural images elicited BoM during the initial V1 response, at a latency of 50ms. This is much earlier than in previous studies that used well controlled, but artificial stimuli to keep the RF stimulus identical between salient and non-salient contour conditions^6, 7^. In these previous studies, the contextual effects on neuronal firing rates were attributed to feedback from higher cortical areas and/or lateral connections within V1, which can inform neurons about information outside the RF. The synaptic and propagation delays associated with these recurrent routes explain why BoM occurs a few tens of ms after the initial V1 response^13^. Our results indicate that the early BoM signals evoked by natural images are not contextual but reflect the tuning of V1 neurons. Indeed, we found that features of object borders differ from those of non-border image regions (Fig. 5c) and that V1 neurons are sensitive to these feature differences (Fig. 5d). On average, the object borders of a particular contrast elicit more activity than non-border image regions with the same contrast. The V1 tuning to object borders is more complex than can be described by Gabor filters^25, 28, 33^ and is presumably related to a sensitivity to higher-order image statistics^34–36^, which also explain the early detection of boundaries in studies using synthetic figure-ground displays^11^ (Fig. 1b).

In addition to their effect on the feedforward response, object boundaries also elicited a contextual influence on V1 activity. When we matched the image elements of object and non-object contours in the RF of V1 neurons, the activity elicited by the object contours was still stronger than that elicited by other, non-object contours (Fig. 6). BoM now occurred at a latency of 81ms, which is 30ms later than the feedforward response and in line with previous studies that used synthetic stimuli to keep the RF content constant and controlled contour salience by the layout of image elements in the RF surround^6, 7^. This additional delay suggests that BoM now depended on feedback from higher areas and/or horizontal connections within V1. It is of interest that these putative feedback signals increased the activity elicited by contours that are predicted by an object’s overall shape. This result is not in accordance with popular “predictive coding” schemes^37^, which suggest that feedback connections should suppress the activity of contours that are predicted by the object’s shape. Instead, we found that object borders increase the neuronal activity in the visual cortex, both during the early and later phases of V1 response.

BoM is presumably related to border-ownership coding, which is expressed by many neurons in V2, V3, V4 and also by some V1 neurons^38–40^. The activity of neurons with border-ownership signals depends on the side of the figural region relative to the border that falls in the RF. For example, if the border is vertical, some neurons prefer that the border is owned by a figure on the left of it, whereas other neurons have the opposite preference. Hence, border-ownership neurons can link the shape of the border to the surface properties of the object’s interior and may therefore play an important role in object recognition. In many situations, the local shape of a border falling in a RF can provide information about the side of the figure^24^. In these situations, neurons express border-ownership early, during the feedforward response. However, if the RF-stimulus is held constant, border-ownership coding occurs after an additional delay^40^. Although BoM reflects extra activity elicited by the object boundaries compared to less-relevant image elements, and thereby differs from border-ownership coding, it seems likely that the two effects are intimately related.

### Neuronal activity elicited by object interiors

Image elements that were part of the interior of objects in the scene elicited more activity than background elements, both in human fMRI and monkey neurophysiology. This finding generalizes previous results on the neuronal mechanism of figure-ground perception to natural images (Fig. 1b,c)^10–12, 16, 41–46^. Studies using synthetic stimuli revealed a number of successive processing phases for the processing of texture defined figure-ground stimuli (Fig. 1b, reviewed in ref.^31^). In V1, the first phase is the arrival of the input from the LGN at a latency of ∼40ms. This is followed at a latency of ∼60ms by boundary enhancement. Boundaries between figure and ground now elicit extra activity in V1, an effect that starts in the superficial layers of cortex^41^. The change in feature values at a boundary between figure and ground can be detected locally (e.g. there is an abrupt change in the texture as in Fig. 1b) and the mechanisms presumably overlap with early BoM (Fig. 7a). In a yet later phase, at a latency of ∼90ms, V1 neurons that represent the figure’s interior enhance their activity. Enhancement in the figure’s interior is a genuine contextual effect, because the properties of the image elements that fall into the RF are often not informative about whether they belong to figure and ground (Fig. 1b,c). In these cases, the information that a RF falls on a figure comes from outside the RF. The relatively long latency of this figure-ground modulation is compatible with recurrent loops that may include horizontal connections within V1 and loops through the higher visual areas. Indeed, if activity in higher areas is blocked, figure-ground modulation in the center of the figure is diminished^47, 48^, implying an import contribution of recurrent routes through higher visual cortical areas^49^. Interestingly, the optogenetic blockade of the late V1 activity phase with figure-ground modulation selectively impairs figure-ground perception, whereas contrast detection is unimpaired^48^.

**Figure 7.**
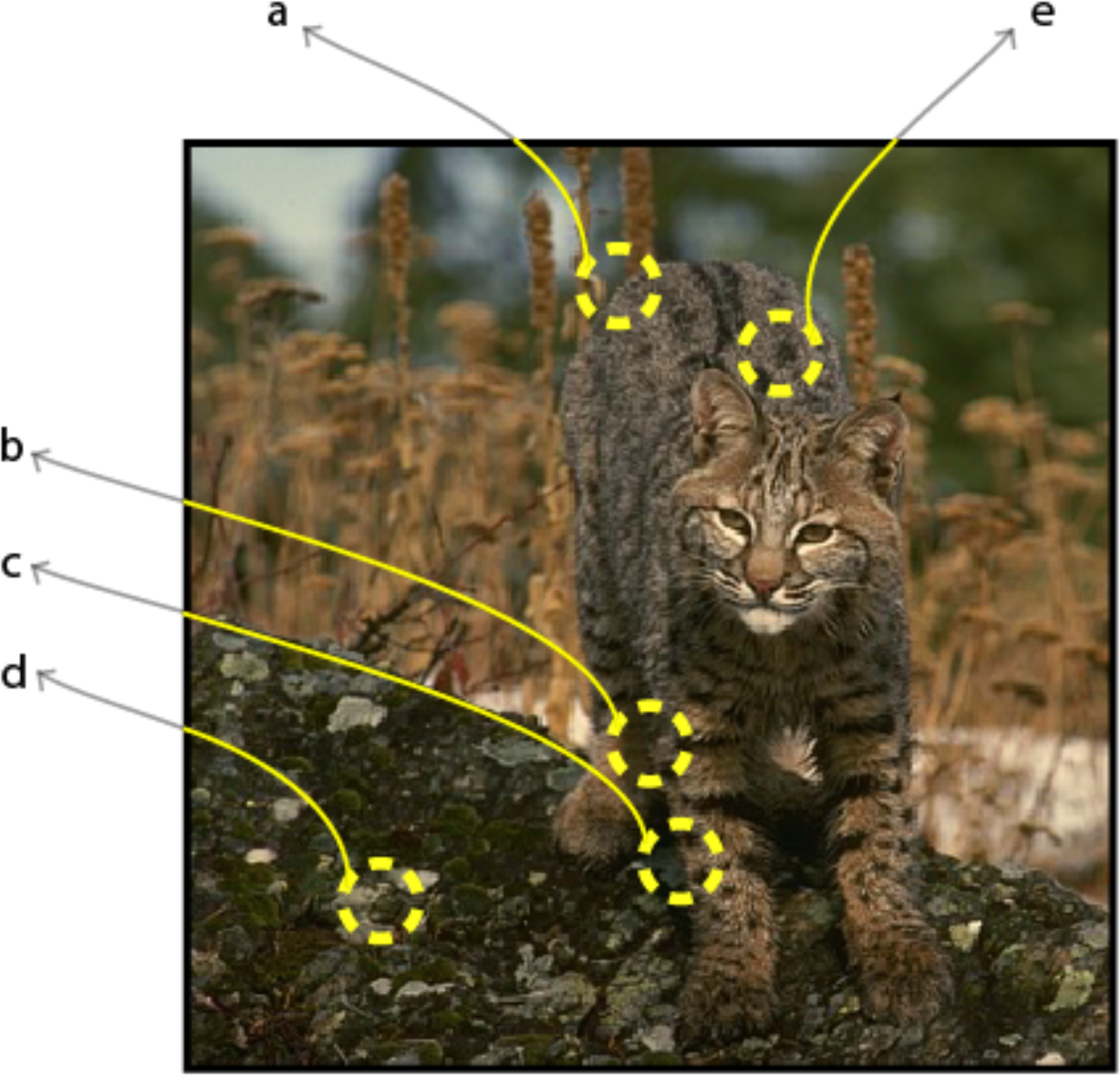
Bottom-up and top-down mechanisms for object detection in natural vision Summary of the results. The local features of contour. **a** suggest that is an object boundary. It can be detected bottom-up by tuning of V1 neurons. Image patches **b** and **c** have similar features but the context indicates that **c** contains an object boundary and **b** does not. Image patches **d** and **e** have similar features, but only **e** is part of the interior of the animal.

The activity of image elements that were part of the interior of objects of natural images was enhanced in V1 at a latency of 78ms, which is 28ms after the visually driven response during the first fixation. In V4 this OBM signal occurred at a latency of 93ms. We also observed systematic differences between the features of object interiors and those in the background, indicating that the tuning of V1 neurons could, in principle, discriminate between features of figure and background, even though the early V1 population response did not exhibit OBM.

### Trans-saccadic integration

Previous studies demonstrated that figure-ground signals for synthetic stimuli can persist across eye movements^17, 18^. As a result, the figure-ground structure that is perceived during one fixation can be quickly reassigned to the appropriate neurons after all RFs shifted across the image due to the saccade. In the present study, OBM occurred sooner after the visually driven response for later fixations than for the first fixation. This result suggests that information about the location of object interiors is indeed carried over to the new fixation^18^, providing insight into the neuronal mechanisms for trans-saccadic integration in natural images^17, 50, 51^. One possible mechanism for the remapping of these response modulations in early visual cortex are neurons in parietal and frontal cortex that remap salient image elements and could provide feedback to lower areas after each saccade^52–54^. Another possible mechanism is provided by neurons that code the position of objects in non-retinotopic, e.g. head-centered coordinates^55, 56^. These cells do not need to update their activity after an eye movement because object position relative to the head is independent of eye position. These neurons could feed the location of objects back to early visual cortex after a coordinate transformation from head to eye centered coordinates, based on the new, post-saccadic eye position. A final source for the early post-saccade OBM are neurons in areas of the temporal stream that code for the overall shape of objects. Many of these neurons are translation invariant, i.e. their activity depends little on the precise location of an object on the retina^57^. These neurons represent object shape^58^ and could provide feedback to boost the activity of neurons in lower areas that represent relevant shape features after a saccade.

## Conclusion

We conclude that the object boundaries and object interiors of natural images increase neuronal activity in the visual cortex. The extra neuronal activity occurs early if the local image elements in the RF have a high degree of “objecthood” and at a later point in time if it depends on contextual information outside the RF. OBM and BoM play an important role in perceptual organization^13^, the process that groups image elements of the same object together and segregates them from other objects and the background by labeling the object features with enhanced neuronal activity^13, 59–63^. The presence of these cortical image parsing signals for natural images suggest that they play a role during each fixation of our everyday vision, opening many avenues for future research.

## Acknowledgments

We thank Kor Brandsma and Anneke Ditewig for biotechnical support. The work was supported by the European Union’s Horizon 2020 and FP7 Research and Innovation Program (Framework Partnership Agreement No. 650003 [HBP FPA]) to PR, the European Union’s Erasmus+ program (grant agreement: 2018-1-IT02-KA103-047276/10) to PP and the Netherlands Organization for Scientific Research (NWO) Vidi (452.08.008) and Vici (016.vici.185.050) to SD.

## Methods

### fMRI experiment with human participants

#### Subjects

Four participants (all male; ages 29-41 years) participated in the fMRI experiment. All participants had normal or corrected-to-normal visual acuity. We obtained informed written consent of the participants and the protocol was approved by the Human Ethics Committee of University Medical Center Utrecht.

#### Stimulus presentation

The visual stimuli were generated in Matlab (Mathworks Inc.) using the PsychToolbox^64, 65^ on a Macintosh MacBook Pro. The stimuli were back-projected on a display inside the MRI bore. The subject viewed the display through mirrors inside the scanner. The size of the display was 15.0×7.9cm with a resolution of 1024×538 pixels. The total distance from the subject’s eyes to the display was 41cm. The stimuli were constrained to a circular area (radius, 5.5°) with the size of the vertical dimension of the screen. The area outside this circle was maintained at a constant mean luminance.

#### Population receptive field (pRF) mapping stimulus

We used bar apertures filled with natural images ^19, 20^ (Fig. S2) to train the pRF-model. The width of the bar subtended 1/4th of the stimulus radius (1.375°). Four bar orientations (0°, 45°, 90° and 135°) and two different step directions for each bar were used, giving a total of 8 bar directions within a given scan. The bar stepped across the stimulus aperture in 20 steps (with a distance of 0.55° and a duration of 1.5 seconds per bar position) so that each pass took 30 seconds. A period of 30 seconds mean-luminance (0% contrast) was presented after every pass. In total there were 4 blocks of mean-luminance during each scan, presented at evenly spaced intervals. The participants performed a fixation dot task to make sure they fixated at the center of the display. A small fixation dot (0.11° radius) was presented in the middle of the stimulus. The fixation dot changed its color from red to green at random time intervals and subjects were instructed to respond to color changes using a button press.

#### Natural images

The natural images came from the BSD ^1, 21^. The original resolution of the images was 321×481 pixels (both landscape and portrait). In the fMRI experiments^20^, we selected a square region of 321×321 pixels from the images and upsampled it to a resolution of 516×516 pixels, which corresponds to a stimulus of 11×11° diameter of visual angle. The images were masked by a circle with a raised cosine faded edge (width of 0.9°), and the areas outside this circle were set to the mean luminance. The images were gamma-linearized and the mean contrast was set to 50%. We used 3 image sets in different scanning runs, each containing 15 different natural images (45 in total) and one full-field binarized bandpass-filtered noise stimulus. Figure S1 shows the image set. A fixation dot was presented at the center of the stimulus. We used the same fixation dot task as for the pRF mapping runs.

#### Functional imaging and processing

The MRI data was acquired with a Philips 7T scanner using a 32-channel head-coil^20^. We scanned the participants with a 2d-echo-planar-imaging sequence with 25 slices oriented perpendicular to the calcarine sulcus with no gap. The following parameters were used; repetition time (TR) = 1500ms, echo time (TE) = 25ms and a flip angle of 80°. The functional resolution was 2×2×2mm and the field of view (FOV) was 190×190×50mm. We used foam padding to minimize head movement. The functional images were corrected for head movement between and within the scans^66^. For computation of the head movement between scans, the first functional volumes for each scan were aligned. Within scan motion correction was then computed by aligning the frames of a scan to the first frame. The duration of the pRF mapping scans was 372 seconds (248 time-frames), of which the first 12 seconds (8 time-frames) were discarded to eliminate start-up magnetization transients. During the three sessions we acquired 6-8 pRF mapping scans in total per subject. To obtain a high signal-to-noise ratio, we averaged across the repeated scans. During the three sessions in which we presented the natural images we acquired 6-7 scans for each of the three stimulus sets. The duration of the scans with the natural images was 432 seconds (288 time-frames). The first 12 seconds (8 time-frames) were discarded to eliminate start-up magnetization transients. The images were presented in a block design. Each image was presented during a 9-second block. Within this block the same image was shown 18 times for a duration of 300ms followed by 200ms mean-luminance. The full-field stimuli were presented with 3 alternating different high-contrast patterns, to obtain a full high-contrast response that is not based upon one specific high-contrast pattern (Fig. S2b). Specifically, the phase of the full-field pattern was randomized on different presentations in order to obtain a response that is not influenced by one specific dartboard pattern. The block in which the stimulus was presented was followed by a 12 second mean-luminance presentation. Four longer blank periods of 33 seconds were also included during the scan.

#### Anatomical imaging and processing

The T1-weighted MRI images were acquired in a separate session using an 8-channel SENSE head-coil. The following parameters were used: TR/TE/flip angle = 9.88/4.59/8. The scans were acquired at a resolution of 0.79×0.80×0.80mm and were resampled to a resolution of 1mm^3^ isotropic. The functional MRI scans were aligned with the anatomical MRI using an automatic alignment technique^66^. From the anatomical MRI, white matter was automatically segmented using the FMRIB’s Software Library (FSL)^67^. After the automatic segmentation it was hand-edited to minimize segmentation errors^68^. The gray matter was grown from the white matter to form a 4mm layer surrounding the white matter. A smoothed 3D cortical surface can be rendered by reconstruction of the cortical surface at the border of the white and gray matter^69^.

#### pRF model-based analysis

The pRF-model was estimated for every cortical location from the measured fMRI signal that was elicited by the pRF mapping bar stimuli (Fig. S2a)^19, 20^. In short, the method estimates the pRF by combining the measured fMRI time-series with the position time course of the visual stimulus. A prediction of the time-series is made by calculating the overlap of the pRF and the stimulus energy (RMS contrast, see below) convolved with the hemodynamic response function (HRF). We estimated the parameters of the HRF that best describes the data of the whole acquired fMRI volume^70^. The optimal parameters of the pRF-model are chosen by minimizing the residual sum of squares between the predicted and the measured time-series. We used the conventional pRF-model, which consists of a circular symmetric Gaussian. This model has four parameters: position (x, y), size (σ) and amplitude (β). For further technical and implementation details see^19^.

#### Regions of interest

We used the pRF-method to estimate position parameters x, and y of the pRF of every voxel. From these values, we derived the polar angle (*atan*(y_0_/x_0_)) and eccentricity (√(x_0_^2^ + y_0_^2^)) values. We drew the borders between visual field maps on the basis^71^ polar angle and eccentricity maps on the inflated cortical surface^69^. We defined visual areas V1, V2, V3, hV4, LO-1/2 and V3-a/b as our regions of interest (ROIs)^71–74^.

#### Analysis of fMRI responses to the natural images

We measured fMRI responses to 45 natural images (Fig. S1) and 3 full-field high contrast stimuli (100% contrast; Figure S2)^20^. We first determined the voxel response amplitudes in %BOLD signal change elicited by each of these images. The voxel responses were calculated using a general linear model (GLM)^75, 76^. To reduce the noise from the individual voxel differences in response amplitudes, we normalized the responses to the voxel’s response to the full-field (100% contrast) stimulus.

To determine the contrast response function (CRF), we only used the voxels with an overall significant response (t-values > 4.0), a pRF eccentricity between 0.5 and 4° and for which the pRF model explained more than 40% of the variance. Based on previous work, for every area we used a threshold for the pRF sizes^19, 70, 77, 78^. In V1 we included pRFs with a value of σ (which determines pRF size) between 0.25° and 0.8°, for V2 between 0.25° and 1.1°, for V3 between 0.25° and 1.75°, for hV4 between 0.45° and 3°, for V3ab between 0.45° and 3.75° and for and LO12 between 0.9° and 5°.

To derive the CRF, we computed the contrast of every natural image within each pRF. The pRF of voxels was modeled as a circular symmetric Gaussian function, described by parameters for position (x_c_, y_c_) and size (σ), giving rise to a Gaussian weighting function w_i_:

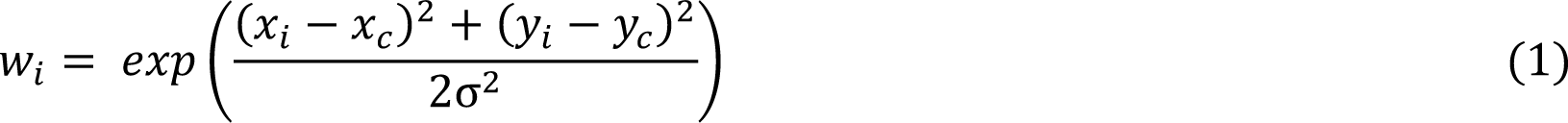

Where x_c_ and y_c_ define the location of the center of the pRF in the visual field, σ determines the size of the pRF and x_i_ and y_i_ define the location of the *i*-th pixel. We computed each voxel’s contrast value to each natural image by calculating the Root-Mean-Squared (RMS) contrast^65, 79^ of the part of the image inside the voxel’s pRF. RMS contrast was defined as the standard deviation of the luminance of the pixels relative to the mean. The RMS-contrast was weighted by the pRF Gaussian function to obtain the local contrast-energy value per pRF:

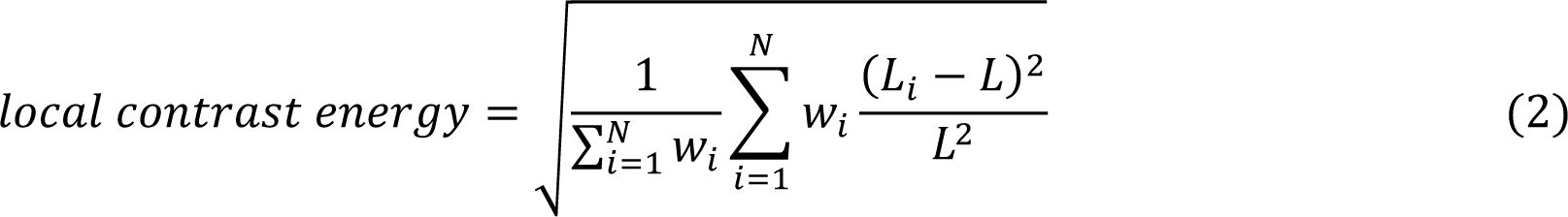

Where *N* is the number of pixels in the stimulus window. *L* is the mean luminance from the pixels inside the spatial window, and L_i_ is the luminance of the *i*-th pixel^2^.

We computed the CRF of voxels areas V1, V2, V3, hV4, LO-1/2 and V3-a/b by measuring the fMRI responses as a function of the contrast inside the pRF. We chose contrast bins such that every bin contained 10% of the voxels and fitted the following equation (modified from ref.^80^):

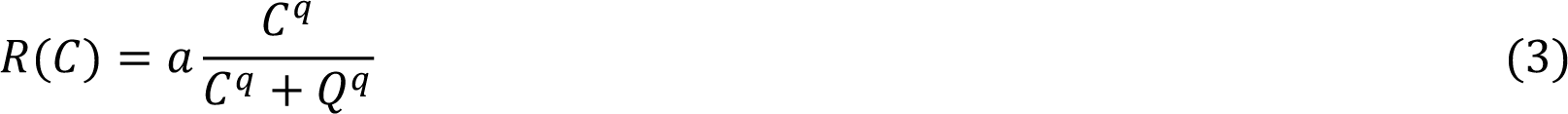

where R is the fMRI response, *C* is the RMS-contrast inside the pRF, *Q* represents the contrast value where the CRF is at half of its maximum response, and *q* determines the slope (*Q* and *q* are free parameters).

#### Quantification of BoM and OBM

The BSD images are annotated by 5-9 human observers who drew lines to identify borders that are important for the scene’s representation^1,21^. We used these measurements to define the perceived boundaries, which are salient boundaries of the scene. Every pixel *i* of the manually labeled images have values for the degree of agreement between observers, *S_i_*, between 0 (not labeled by any observer) and 1 (labeled by all observers). The border-salience in the pRF is calculated as a weighted sum across pixels:

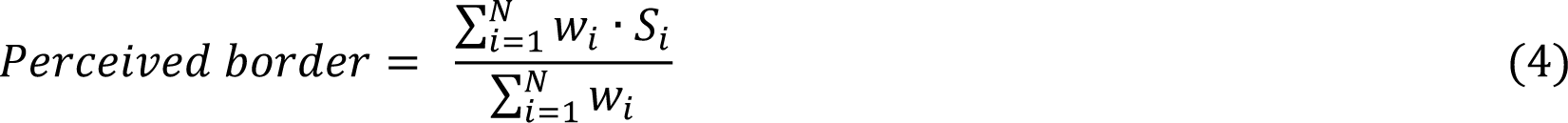

Here *w_i_* are the weights of the RF estimate (equation 1) and *N* is the total number of pixels in the RF. We used the same method to quantify the degree to which a pixel was part of an object or the background (Fig. 1e). Pixels that were as part of an object, had a value of 1 and pixels that were part of the background had a value of 0. We selected a segmentation covering the objects of a scene from one of the BSD subjects, and then considered everything else as background^5^. We excluded 3 of the 45 images in the OBM analysis because the object in the image almost filled the entire scene. We split the voxels based on objecthood values inside the pRFs. We included the lowest 25 percent responses as non-perceived borders/background and the highest 25 percent as perceived-borders/object-interior and computed the CRFs within these voxel classes.

## Statistics

We used a bootstrapping procedure to determine the significance of differences in CRFs between conditions. We sampled the images with replacement 1000 times, fit the CRF for the two simulated conditions and computed the mean difference. We derived the p-value from this null distribution.

### Electrophysiological experiments in monkeys

#### Training of the monkeys

All procedures complied with the NIH Guide for Care and Use of Laboratory Animals and were approved by the institutional animal care and use committee of the Royal Netherlands Academy of Arts and Sciences. Two macaque monkeys (males, 7 and 13 years old) participated in the electrophysiological experiments. They were socially housed in stable pairs in a specialized primate facility with natural daylight, controlled humidity and temperature. The home-cage was a large floor-to-ceiling cage which allowed natural climbing and swinging behavior. The cage had a solid floor, covered with sawdust and was enriched with toys and foraging items. Their diet consisted of monkey chow supplemented with fresh fruit. Their access to fluid was controlled, according to a carefully designed regime for fluid uptake. During weekdays the animals received water or diluted fruit juice in the experimental set-up upon correctly performed trials. We ensured that the animals drank sufficient fluid in the set-up and supplemented extra fluid after the recording session if they did not drink enough. On days of the weekend, they received at least 700ml water in the home-cage in a drinking bottle. The animals were regularly checked by veterinary staff and animal caretakers and their weight and general appearance were recorded daily in an electronic logbook during fluid-control periods.

#### Surgical details

We implanted both monkeys with a titanium head-post (Crist instruments) under aseptic conditions and general anesthesia as reported previously^81^. The monkeys were trained to direct their gaze to a 0.5° diameter fixation dot and hold their eyes within a fixation window (1.1° diameter). They then underwent a second operation to implant 5×5 arrays of micro-electrodes (Utah-probes, Blackrock Microsystems) over opercular V1 and V4. The inter-electrode spacing of the arrays was 400μm. We obtained good signals from 4 V1 arrays in each monkey and from 2 V4 arrays in monkey B^11^.

#### Electrophysiology

We recorded neuronal activity of 192 recording sites in V1 (96 in Monkey M and 96 in Monkey B) and 48 V4 recording sites in monkey B. We recorded the envelope of multi-unit activity by digitizing the signal referenced to a subdural electrode at 24.4kHz. The signal was band-pass filtered (2nd order Butterworth filter, 500Hz-5KHz) to isolate high-frequency (spiking) activity. This signal was rectified (negative becomes positive) and low-pass filtered (corner frequency = 200Hz) to produce the envelope of the high-frequency activity, which we refer to as MUA^82^. The MUA signal reflects the population spiking of neurons within 100-150μm of the electrode and the population responses are very similar to those obtained by pooling across single units^82–85^.

#### Receptive Field Mapping

We mapped the RF of each recording site in V1 using a drifting luminance-defined bar that moved in one of four directions. The response to each direction was fitted with a Gaussian function. The borders of the RF were then calculated as described previously^82^. The signal-to-noise ratio (SNR_RF_) of the response was taken as the peak of the Gaussian divided by the standard deviation of the pre-trial baseline response. We only included recording sites in the analyses with a reliable visual response (i.e., the responses to all four bar directions had an SNR_RF_ of at least 1). The median V1 RF size, taken as the square-root of the area, was 1.8° (range 0.4° to 8.2°) and the median eccentricity of the RFs was 2.4° (range 0.6° to 12.9°). We mapped V4 RFs by presenting white dots (0.5°, luminance 82 cd×m^−2^) on a gray background (luminance 14 cd×m^−2^) at different positions of a grid (0.5° spacing). The hotspot of the V4 RF was defined as the position with the maximum response (median eccentricity 4.04°, range 0.79°–7.43°) and the RF borders as the locations where activity fell below 50% of the maximum^86^. Using this criterion, the median V4 RF size was 4.5° (range 2.6°– 6.0°).

#### Stimulus presentation

In the experiments with monkeys, stimuli were presented on a CRT monitor at a refresh rate of 60Hz and resolution of 1024×768 pixels viewed from a distance of 46cm. The monitor had a width of 40cm, yielding a field-of-view of 41.6 x 31.2°. All stimuli were generated in Matlab using the COGENT graphics toolbox (developed by John Romaya at the LON at the Wellcome Department of Imaging Neuroscience). The eye position was recorded using a digital camera (Thomas recordings, 250Hz frame-rate).

### Selection of recording sites and inclusion of data

To normalize MUA, we first subtracted the mean activity in the pre-trial period in which the animal was fixating (200 to 0ms relative to stimulus onset) and divided by the maximum smoothed (26ms Gaussian kernel) peak response (0-150ms after stimulus onset). In the experiment with multiple saccades, each trial contained multiple fixations and neuronal activity was normalized to the peak response elicited by stimulus onset during the first fixation. The data are therefore in normalized units, where e.g. a value of 0.1 indicates 10% of the maximal MUA onset response. We only included recording sites on days with a sufficient signal-to-noise ratio (SNR_DAY_). SNR_DAY_ was estimated by dividing the maximum of the initial peak response by the standard deviation of the baseline activity across trials. When the SNR_DAY_ of a recording site was smaller than 2 on particular day, we removed that session from the analysis of that recording site. To test for statistical differences between conditions and to compute the CRFs, MUA activity was generally averaged in a 0-300ms time window. Analyses with different time-windows have been specified in the main text.

### Analyses of latency

To compute the latency of neural responses a function was fitted to the time-course of interest (i.e. the difference between object borders and non-border image regions or the difference between the object interior and background)^11, 12, 23^. The function was derived from the assumptions that the onset of the response has a Gaussian distribution and that a fraction of the response dissipates exponentially which yields the following equation:

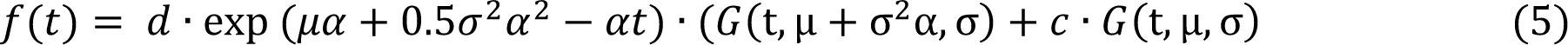

Where G(t, µ, σ) is a cumulative Gaussian density with mean μ and standard deviation σ, α^−1^ is the time constant of the dissipation, and c and d represent the contribution the non-dissipating and dissipating components, respectively. The latency was defined as the point at which the fitted function reached 33% of its maximum. To compare the latency of the BoM and OBM between fixation 1 and fixations 2-6, we first subtracted from the OBM latency for each recording site from the latency of visually driven response and performed a Wilcoxon signed rank test (Fig. S7). The latency of visually driven response was computed as the difference between the response elicited by images with the highest and lowest contrast levels in the RF.

### Natural images presented in the electrophysiological experiments

Four BSD images from the fMRI experiment were used in the electrophysiological experiments in which the monkeys made saccades (11.6° radius visual angle; Figure S1). At the start of the trial the screen was gray (26.8 cd×m^−2^) with a red fixation point with a position that was randomly selected from uniformly spaced grid (with ∼500 positions) covering the circular aperture of the image. The image appeared once the monkey had maintained fixation for 300ms (fixation 1). After an additional 400ms, the first fixation point disappeared and another fixation point appeared, at a position sampled from the same grid. The monkey made a saccade to the new fixation point and maintained fixation for an additional 400ms. This fixation-saccade procedure was repeated five times (fixations 2-6). Reward was delivered after every correct fixation, with an extra amount at the end of the trial, i.e. after the 6^th^ correct fixation. Aborted trials (i.e., when the monkeys did not maintain fixation for 400ms or did not perform a saccade within 700ms) were repeated at the end. The same image was presented in multiple recording days until data for five repetitions of each grid point for every fixation number was collected. We included data from all correct fixations (e.g., if the trial was interrupted after five fixations, we included the first four). Between the trials, the monkeys occasionally also fixated on parts of the image for longer than 300ms, and we also included these spontaneous fixations in the analysis. We collected a total of 11,783 correct trials for monkey M and 13,373 for monkey B, for a total of 50,849 fixations analyzed for monkey M and 60,211 for monkey B.

### Data analysis

We determined the coordinates of the RF on the image for every fixation and analyzed the data from the first fixation and later fixations separately. We computed contrast, BoM and OBM in the RF, as described above. To quantify the independent influence of object borders, object interiors and contrast, we carried out a variance partitioning analysis^87^. For each recording site, we determined how much variance (R^2^) was explained by RMS contrast, object borders and object interiors with independent linear regressions, and by combinations of the three predictors in multiple linear regressions. For example, the independent fraction of explained variance (FEV) for the contrast predictor was computed for every recording site as follows:

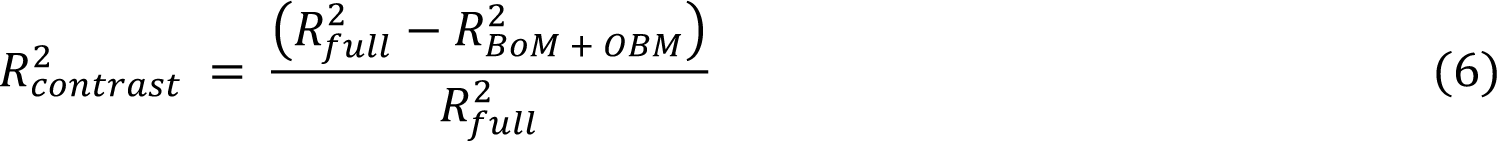

where R^2^_full_ is the variance explained by the full model, including all the predictors, while R^2^_BOM + OBM_ is the variance explained by the model including BoM and OBM as predictors while leaving contrast out. Similar equations were derived for the FEV accounted for by BoM and OBM.

The explained variance estimates were then averaged across recording sites. The full model explained 6.4% of the variance in V1 for fixation 1 (mean across recording sites), 4.3% for fixations 2-6, 3.2% in V4 during fixation 1 and 2.5% for fixations 2-6. The FEV values for each area and condition presented in the main text were normalized to these values (see equation 6).

### RF models and the prediction of perceived borders

We determined the selectivity of the neurons at a recording site (time-window 25-75ms), according to previous studies^25–28^ which established a mapping between an artificial neural network (ANN) and neuronal tuning (Fig. S9).

We extracted the activity of units of VGG-19’s layer conv3_1 (state of the art in predicting V1 responses to natural images^28, 29^) and followed the approach of ref.^28^ with two modifications. We used a two-step mapping^26, 27^, described by following the equation:

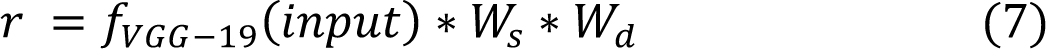

where r is the predicted response of V1 recording site, f_VGG-19_(input) is the output of VGG-19’s conv3_1 to our stimulus set (i.e. input), and W_s_ and W_d_ are two sets of weights defining spatial and feature selectivity, respectively. The spatial mask (W_s_, initialized as the 2D Gaussian RF estimate) approximates the RF and a weighted sum of the nodes in the ANN (W_d_) approximates the feature selectivity of the recorded neurons^26^. We trained the model to optimize W_s_ and W_d_ to predict V1 responses to the training set (i.e., input in eq. 7). The activity of V1 recording sites depends on a small and localized portion of the input and we therefore cropped the RF models around the most active pixels (3 SD or more away from the global mean) following the procedure in ref.^25^. To visualize tuning (Fig. 5b,c), we kept W_s_ and W_d_ constant and varied input to maximize the response of the model for a particular V1 recording site. We used cross-validation to assess the quality of the fit as in ref.^25^. Specifically, we used 5,000 trials for training and 100 trials for cross-validation. We trained the model for 300 epochs with a batch size of 256, using 10% of the training set for validation.

To estimate how well the V1 RF models could detect perceived borders (F-stat metric; Fig. 5b), we first convolved the RF models with unseen images from the BSD test set (100 grayscale images), and matched them with the annotated versions^21^. For simplicity, we defined the border detection performance (BoP) as the F-measure, employed by the authors of the BSD for benchmark evaluation, which we computed using the MATLAB code associated with the dataset: (https://www2.eecs.berkeley.edu/Research/Projects/CS/vision/bsds/code/)20. We estimated the chance-level performance in border-detection with a (null) permutation distribution (horizontal dashed line in Fig. 5c; 97.5% level the distribution), shuffling the class labels (after 72 iterations, one for each recording site included in this analysis. A generalized Pareto distribution was fit to the tail of the permutation distribution^88^. The models were implemented using custom Python code using NumPy^89^, SciPy (SciPy.org), Tensorflow 1.5^90^ and with modules from https://github.com/dicarlolab/npc26 and https://github.com/sacadena/Cadena2019PlosCB28.

### Perceived borders and object interior detection of a population of V1 neurons

To examine the strength of BoM and OBM signals across a larger population of V1 recording sites (Fig. 5d), we trained SVMs to discriminate between object borders and non-border image regions based on the activity of 19 recording sites with RFs smaller than 1.5°. We also trained them to distinguish between object interiors and the background. We used 2 of the 4 images for training and the other two for cross-validation.

### Neuronal activity profiles across the images

To examine the overall activity level elicited by the images (Fig. 3b), we multiplied the activity by 2d-Gaussian approximation of the RFs, weighted by sampling of the visual space caused by the overall pattern of fixations^5,20, 91–95^ at 7 time points (from 0 to 300ms in 50ms steps) and activity was averaged within a 25ms window centered on each time-point.

### Statistics

We compared differences between the CRFs between object borders and non-border image regions and between object interiors and the background using a bootstrapping procedure (1,000 iterations), as described for the fMRI data above. To test for differences in the median latencies of BoM and OBM between regions and conditions, we used a signed-rank Wilcoxon test across recording sites. The significance of the Pearson’s correlation between BoM at the peak of response and the segmentation performance across V1 recordings sites (Fig. 5b) was assessed with a t-test.

### Isolated patches experiment

To test whether isolated image patches from the BSD that were either centered on object borders or not elicited a different level of V1 activity, we carried out an additional experiment in monkey B (50 recording sites, Fig. 5e-g). We chose three V1 recording arrays and centered 100 patches of the image from the BSD that contained object contours and 100 patches that did not contain object contours on the RFs. These patches were automatically selected so that the RMS contrast was the same (70±1%) and the size matched to the median RF of the recording sites of the array (0.9° − 2.0°). The patches were presented on a grey background (26.8 cd×m^−2^) while the monkey maintained gaze on a red fixation point for 300ms. We repeated each stimulus five times and collected a total of 3,000 trials (1,000 trials per array). We tested the significance of the difference in the activity elicited by isolated object and non-object contour patches at the peak of the response (25-75ms) with a Wilcoxon signed rank test across recording sites.

### Contextual BoM experiment

To examine differences in activity elicited by object and non-object contours when the stimulus in the RF was held constant (Figure 6) we selected twelve images from the BSD, which were cropped and upsampled to 512 x 512 pixels (23.2° x 23.2°). We ensured that the portion of the image covered by the RF of each recording site and its surround were exactly the same across conditions (same size and content, Fig. 6), so that border salience only depended on information outside the neurons’ RF. We used a 2×2 design. The first factor was whether the image element in the RF fell on an object border (Fig. 6a). The second factor was whether we presented the original image or a scrambled version (also known as *metamer*). To this aim, we created three further stimuli from each image. First, we copied a circular patch (80 pixels in diameter, 3.7°) from an object contour location onto a example images). The border of this circular patch was smoothed to blend it in at the new location. We created two metamers using the algorithm of ref.^96^, with Matlab code provided by the authors (https://github.com/freeman-lab/metamers). The two metamers were constructed so that either the object- or non-object contour was kept intact, with a smooth transition to the surround.

Trials started with a red fixation point and the stimulus appeared after 300ms of fixation. The monkeys maintained fixation for an additional 400ms after stimulus onset (Fig. 6b). We ensured that the RFs of V1 recording sites were centered on the image patch, which was identical in the four conditions. The order of the conditions was randomized across trials and aborted trials (when the monkeys broke fixation) were repeated at the end. We collected a total of 8,094 trials in monkey M and 9,111 in monkey B.

We tested the significance of the BoM in a window from 0-300ms after stimulus onset (subtracting spontaneous activity, −100-0ms) with a Wilcoxon signed rank test across recording sites. We also used a repeated-measures two-way ANOVA across recording sites, with object/non-object contour and scrambled/not scrambled as factors.

## Data availability

Data will be available upon publication of the paper.

## Code availability

Custom code will be available upon publication of the paper.

