## Supplementary information for "The influence of objecthood on the representation of natural images in the visual cortex"

*Testing measures of stimulus energy other than RMS Contrast*

There are multiple methods to estimate contrast. In addition to the local RMS contrast, we tested several other methods to predict V1 activity during Fixation 1 (comparable results were obtained also for Fixations 2-6 – not shown). The first three methods were canonical V1 predictors, complex cell energy ($E_{CC}$), cross-orientation energy ($E_{CO}$) and surround suppression energy ($E_{SS}$, Fig. S8a, pink)[97] for patches of the natural images falling into the RF. We evaluated the quality of the predictions by computing the correlation with the V1 activity level elicited by these image patches (Fig. S8). $E_{CC}$ was computed as:

$$E_{CC} = \sqrt{E^2_{x,y,F,\theta,\Phi=0} + E^2_{x,y,F,\theta,\Phi=90}}$$

where $E$ is the dot product of the stimulus with a 2D Gabor with coordinates $x$ and $y$, $F$ is spatial frequency, $\theta$ orientation and $\Phi$ phase. $F$ and $\theta$ are free parameters of the model and were optimized for every recording site before we calculated the correlation coefficient.

Cross-orientation inhibition (CO) was computed as:

$$E_{CO} = \left(\frac{E_{CC}}{\alpha + \beta E_{CO}}\right)$$

where $E_{CO}$ is the sum of complex cell energy computed at 4 orientations ($\theta$ = 0, 45, 90, 135 – the other parameters are the same as for $E_{CC}$) and $\alpha$ and $\beta$ are free parameters. $E_{SS}$ was computed as

$$E_{SS} = \left(\frac{E_{CC}}{\alpha + \beta E_{CO} + \gamma E_{CS}}\right)^n$$

$E_{CS}$ is the sum of complex cell energy computed using the same parameters as $E_{CO}$ but centered radially at the distance of 2 RF radius from the RF center. $\gamma$ and $n$ are the free parameters. Each free parameter was estimated in the simpler models and carried over to the more complex models. $F$ and $\theta$ were estimated for $E_{CC}$ and then used estimate $E_{CO}$, where the new free parameters $\alpha$ and $\beta$ where estimated. Similarly, only $\gamma$ and $n$ were estimated for $E_{SS}$. We used Matlab's *lsqcurvefit* function to compute the fits.

We tested three further predictors for the level of neuronal activity (Fig. S8a, light blue), which have been proposed in previous neuroimaging and behavioral studies. Specifically, we chose the compressive spatial summation model (code from the authors: http://kendrickkay.net/socmodel/)[98], local RMS contrast weighted by a Difference of Gaussians function (DoG)[77], and Sobel magnitude. We additionally tested four models sensitive to texture-homogeneity (Fig. S8a, blue): the H-image model[99], the Mixture of Gaussians Mixture Scales model (MGMS[100]; eq. S8 in ref.[101], code from the authors: https://portal.nersc.gov/project/crcns/download/mgsm-1), a combination of local RMS Contrast and MGMS – computed as:

$$Resp_{MGMS+RMS\ Contrast} = \frac{local\ contrast}{\alpha + w_i * S_{MGSM}}$$

The local contrast was computed as in equation 1, $\alpha$ was set to 1.7 to allow both suppressive and facilitatory modulations, $w_i$ is the Gaussian weighting function described in equation 2 and $S_{MGMS}$ is the local homogeneity (eq. S8 in ref.[101]). The last model was the gPb contour detector[21] from a state-of-the-art hand-crafted segmentation model.

The correlation coefficient for RMS Contrast was larger than or equal to those for the other models, with the exception of the gPb model. We preferred RMS Contrast because it is simpler and has no parameters. BoM occurred irrespective of the precise choice of the model.

The most advanced model was based on a convolutional neural network (VGG-19, third convolutional layer 'conv3_1', as implemented in *MatConvNet*: http://www.vlfeat.org/matconvnet/), which provides state-of-the-art prediction of V1 responses[28]. The output of each node of convolutional layer 3_1 was weighted by the RF of each recording site in each trial, and we used the layer 3_1 activity vector to predict the neuronal response. The (cross-validated) correlation coefficient of this model was higher than that of all the other models, including RMS Contrast (Fig S8a, gray). However, we also observed BoM when we modeled the CRF using this ANN model, just as was the case when we used RMS Contrast (Fig. S8b).

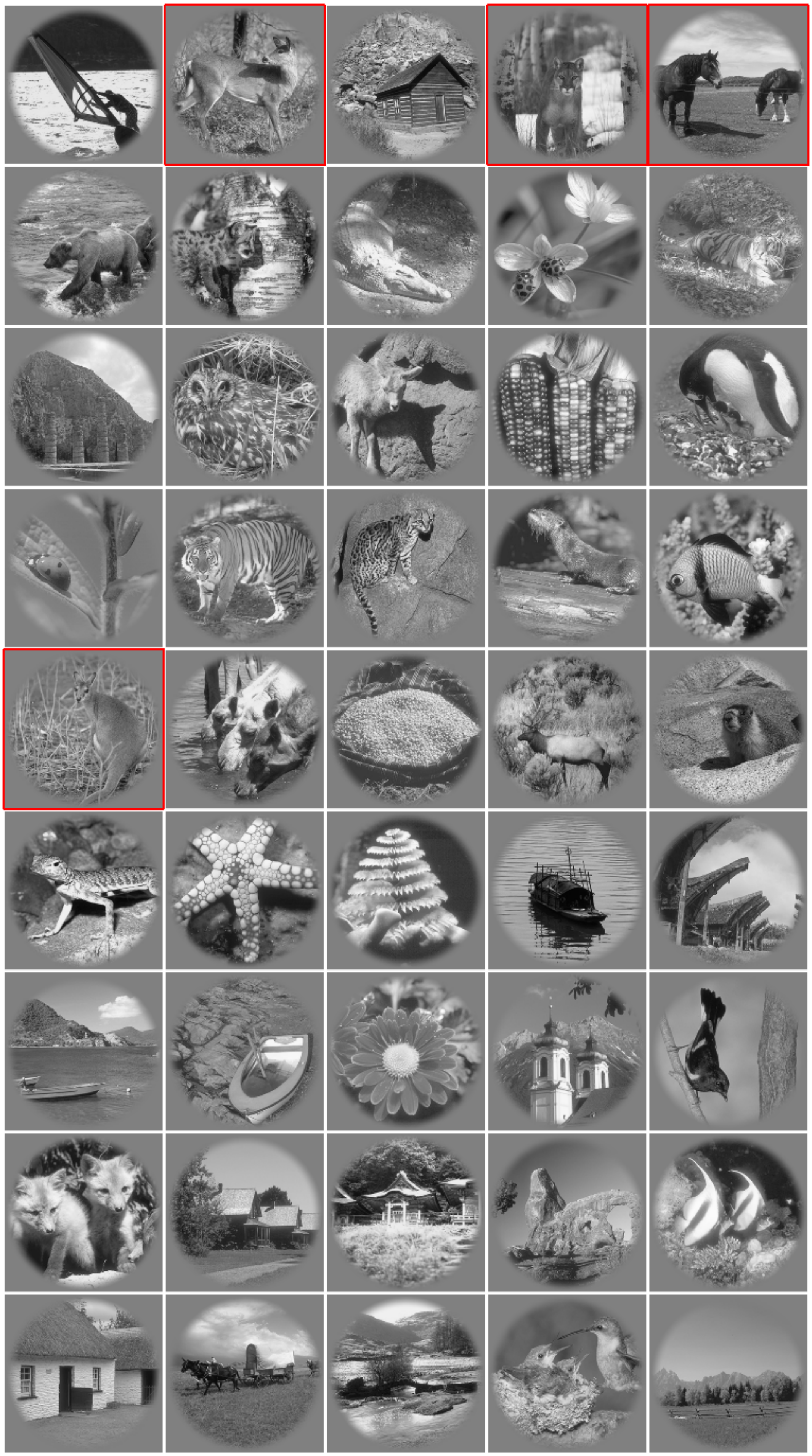

**Figure S1. Natural images.**

Stimuli from the BSD that were used in the fMRI experiments with humans. Red outlines: subset of stimuli that were used in the experiments with monkeys.

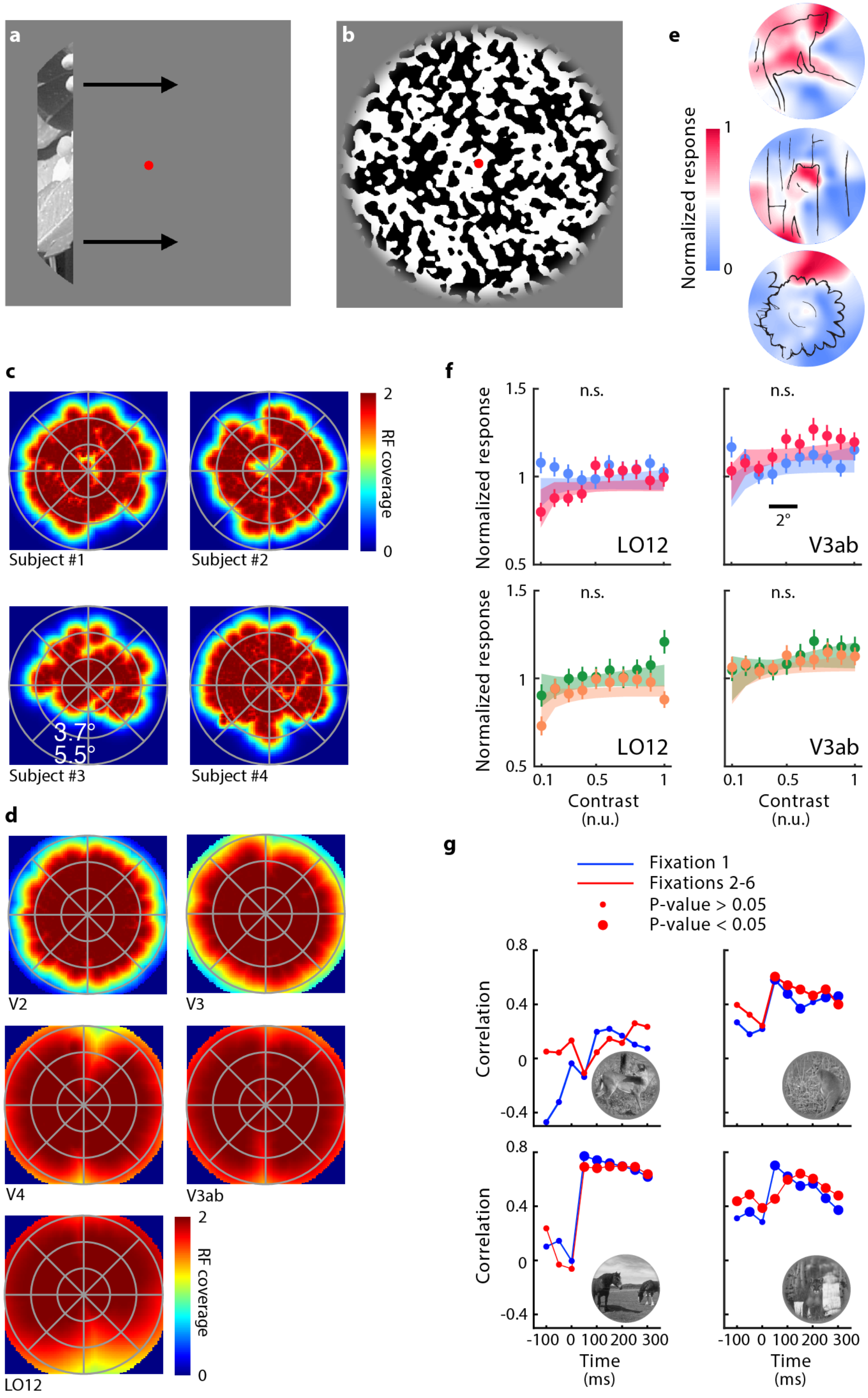


**Figure S2. pRF mapping and CRF in LO12 and V3ab**

**a**, The pRF mapping stimulus of the fMRI experiment consisted of a bar stepping through the visual field in 8 different directions. **b**, We used the response evoked by a full-field stimulus (100% contrast) to normalize the voxels' responses. **c**, V1 pRFs coverage plots for individual subjects. Color indicates the degree of coverage of the visual field with pRFs (maximum value is clipped at 2 pRFs covering a point in space). **d**, Coverage in other cortical areas, averaged across subjects. **e** V1 fMRI responses elicited by three example images. For the upper image, the activity is focused on object boundaries, for the middle image the relation between activity and the boundaries is less pronounced and it is absent for the bottom image. **f**, CRFs in LO12 and V3ab (top: influence of object-borders; bottom: influence of objecthood). **g**, Correlations between the fMRI response in V1 in humans and the spiking activity recorded from V1 neurons in monkeys viewing the same four natural images. For this comparison, we projected both fMRI and electrophysiological responses in the visual space. In monkeys, we binned the response over the natural images across time (centered every 50ms, from -100ms to 300ms after the stimulus onset) separately for both fixation 1 (blue) and fixations 2-6 (red). Correlation coefficients were tested for significance using a permutation test (1,000 iterations by using the fMRI responses to the different images, with random spatial rotations). The size of the dots indicates significance (big dots are significant, small dots not).

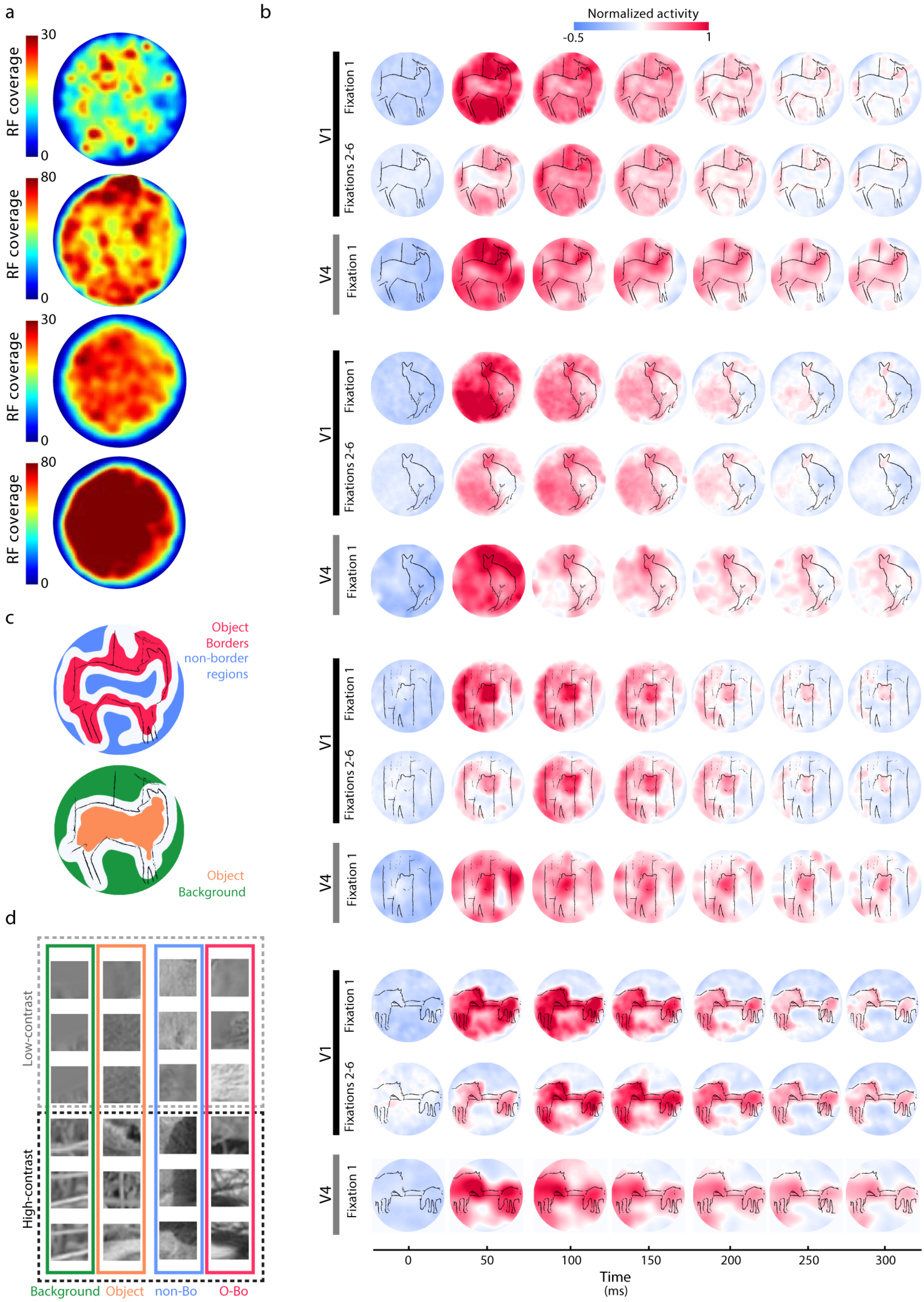

a
RF coverage
30
0
80
0
30
0
80
0
b
Normalized activity
-0.5
1
V1
Fixation 1
Fixations 2-6
V4
Fixation 1
c
Object
Borders
non-border
regions
Object
Background
d
Low-contrast
High-contrast
Background
Object
non-Bo
O-Bo
0
50
100
150
200
250
300
Time
(ms)

**Figure S3. MUA in areas V1 and V4 evoked by natural scenes.**

**a**, RF coverage for an example image (from top: V1 Fixation 1: V1 Fixations 2-6; V4 Fixation 1; V4 Fixations 2-6). Color indicates the degree of coverage by RFs. **b**, V1 and V4 MUA elicited by the images across time. The first two rows per image represent V1 activity elicited during fixation 1 and fixations 2-6, respectively. The third row illustrates V4 activity during fixation 1. Note that V4 activity during fixations 2-6 is illustrated in Fig. 2b. **c**, Assignment of image regions to boundary, interior and background for an example image and recording site. **top,** The area of the image falling on the RF that counted as object border (non-border image region) is shown in red (blue). White regions are ambiguous and excluded from the analysis. **bottom,** Image regions used to determine the influence of objecthood. The object-interior is shown in orange and the background in green. **d**, Image regions falling in the RF of an example V1 recording site at the lowest and highest contrast for object-borders (red) and non-border image regions (blue) and for object-interior (orange) and background (green). The difference in local contrast is evident, but objecthood differences are not as clear, because they depend on the context provided by image regions outside the RF.

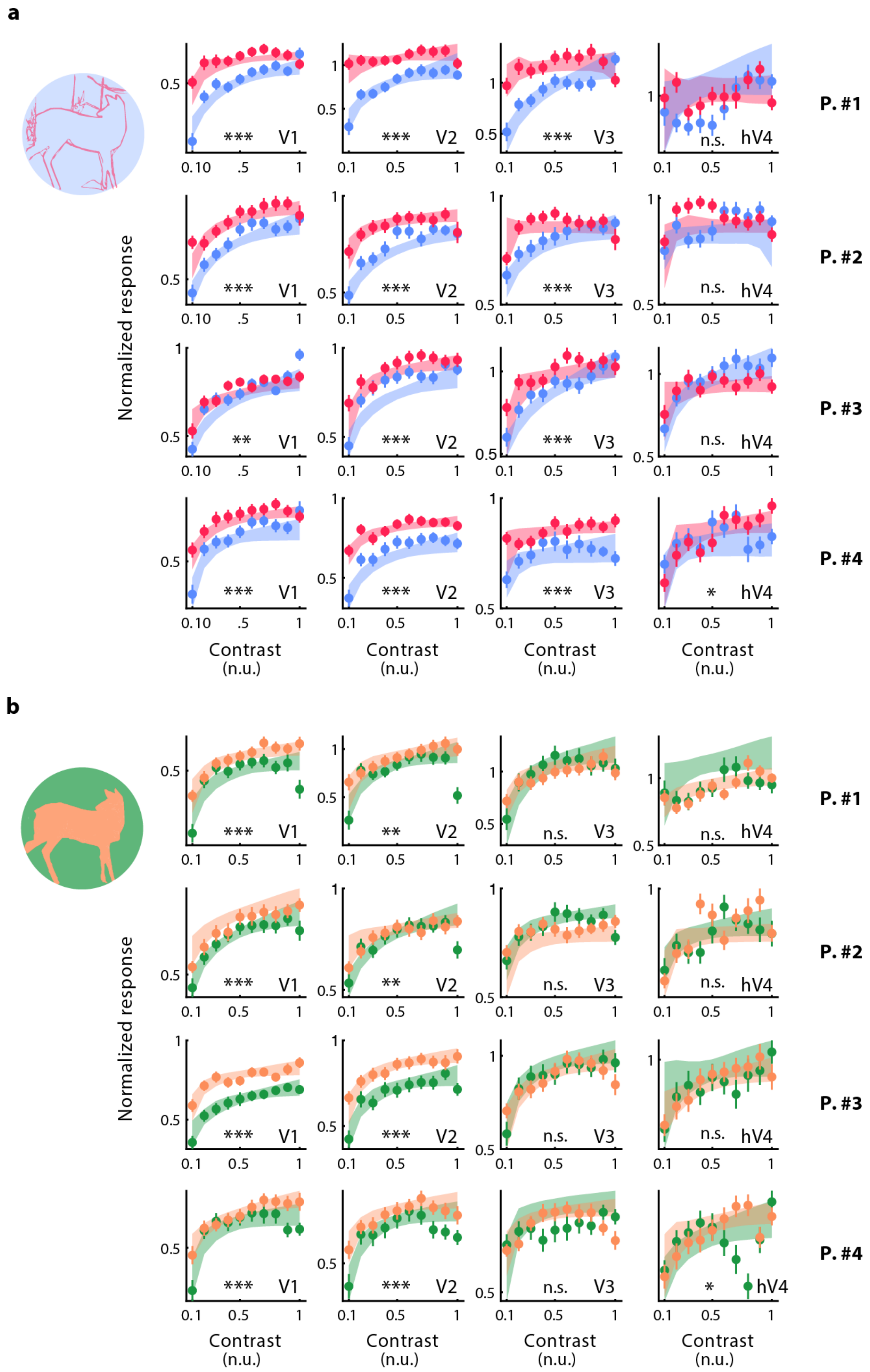


**Figure S4. BoM and OBM in V1 to V4 of the four fMRI participants**

**a**, Same as Figure 2a: BoM in V1 to V4 (columns) of individual participants (rows).

**b**, Same as Figure 2b: OBM in V1 to V4 (columns) of individual participants (rows) (*: $p < 0.05$; **: $p < 0.01$; ***: $p < 0.001$, bootstrap test).

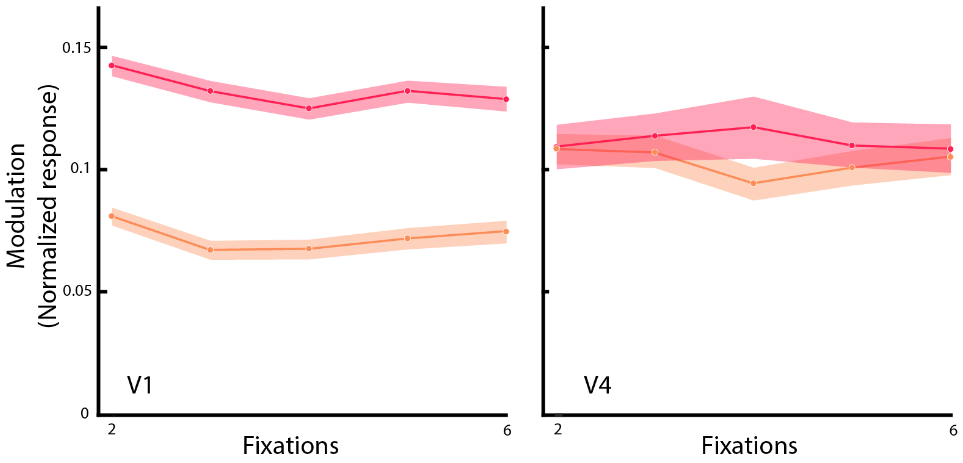


**Figure S5. Consistency of BoM and OBM across fixations 2-6.**

BoM (red) and OBM (orange) in V1 (left) and V4 (right) for fixations 2 to 6 (shaded areas: 95% bootstrapped CI). All values are significantly larger than zero ($p < 0.001$, bootstrap test).

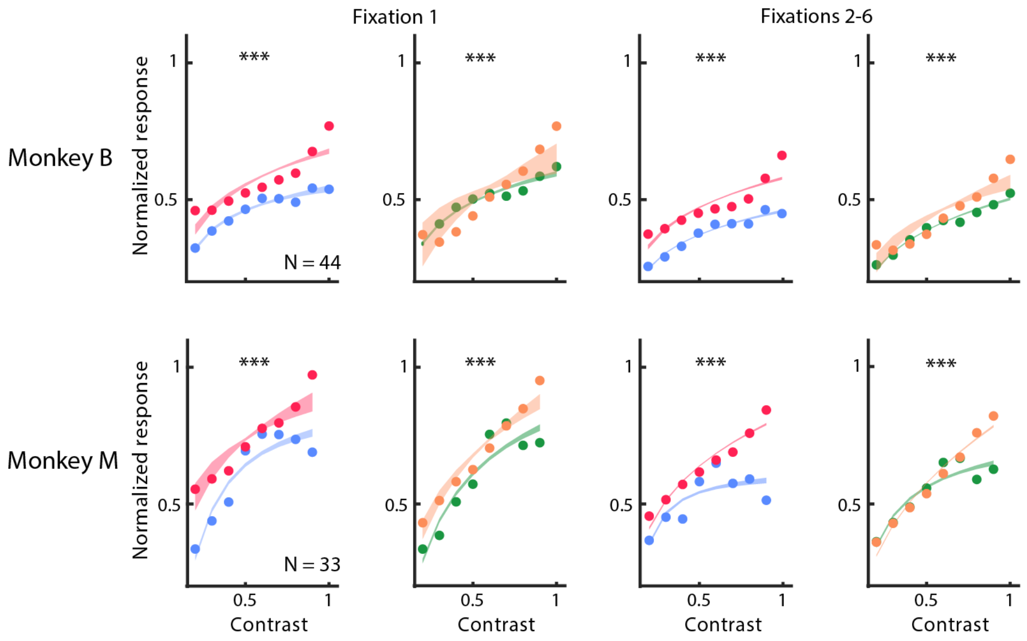


**Figure S6. BoM and OBM in V1 of the two monkeys.**

BoM (difference between red and blue curves) and OBM (orange vs. green curves) for the two monkeys. BoM and OBM are significant in all conditions in both animals (***, $p<0.001$).

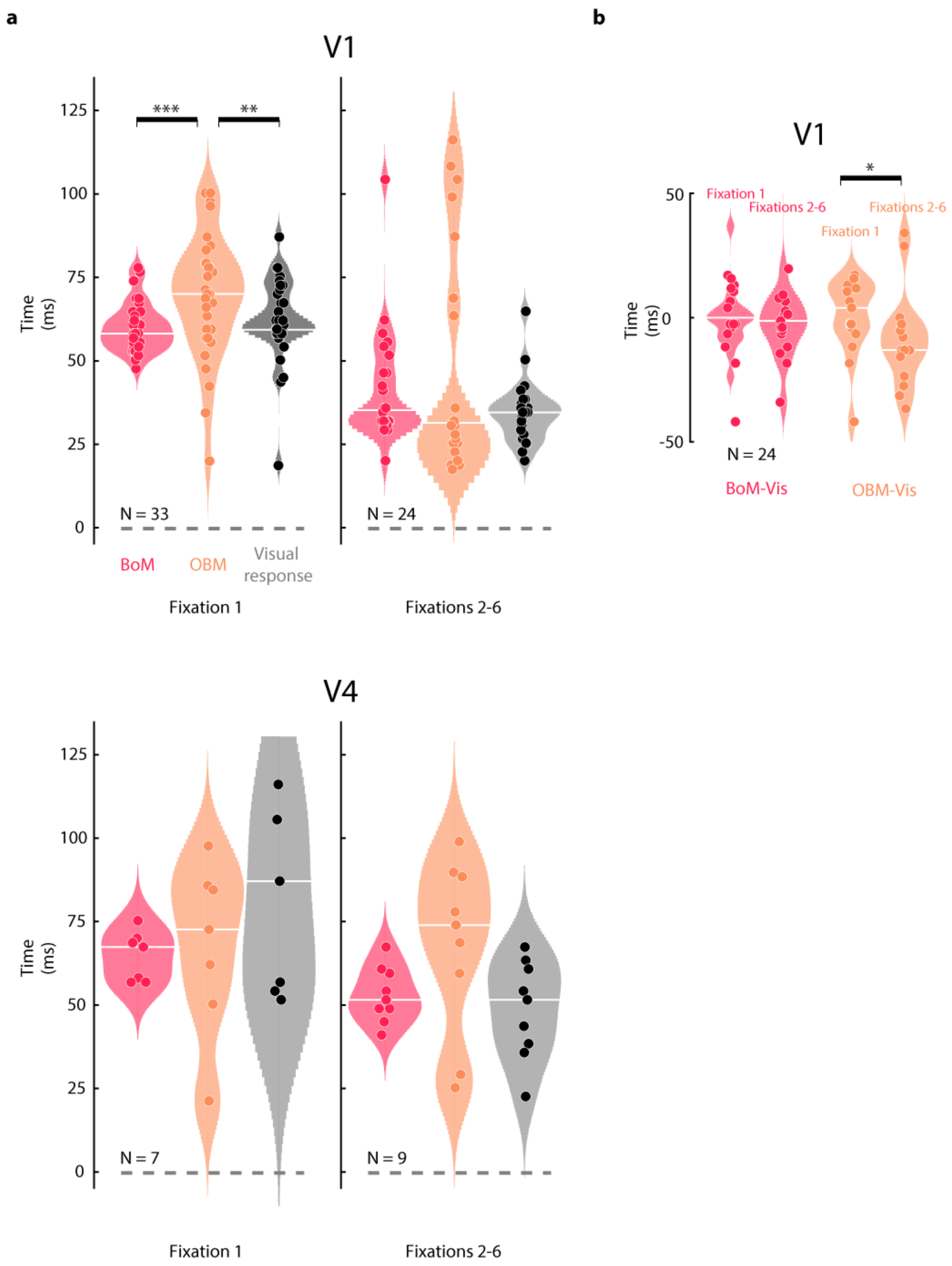


**Figure S7. Distribution of latencies of the visual response, BoM and OBM across recording sites.**

**a**, Latency distributions of the visual response (black), BoM (red) and OBM (orange) across V1 sites with significant modulation. Horizontal white lines indicate the median latency. During the first fixation, OBM is significantly later than both the visual responses and BoM. **b**, To compare latencies across sites in V1 between fixation 1 and fixations 2-6, we first computed the difference between the BoM latency and the visual latency ($Lat_{BOM\text{-}Vis}$) and the same for OBM ($Lat_{OBM\text{-}Vis}$). $Lat_{OBM\text{-}Vis}$ was significantly shorter during fixations 2-6 than during fixation 1. This comparison did not significance in V4, but this is presumably due to the lower number of V4 recording sites (N=7). (*: $p < 0.05$; **: $p < 0.01$; ***: $p < 0.001$, Wilcoxon signed rank test).

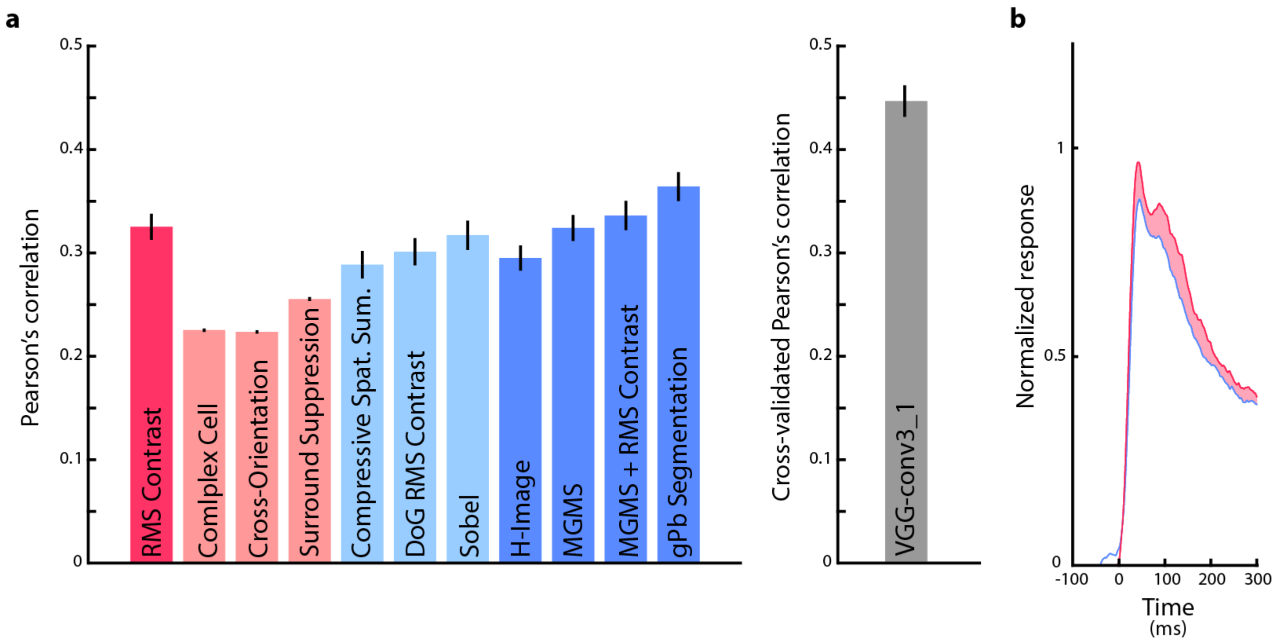


**Figure S8. Predictors of the magnitude of the V1 responses.**

**a**, Correlation between predictors of V1 activity and the V1 response elicited by natural images (see supplementary text above for a description of all predictors). **b**, Time-course of spiking activity at the population level in V1 (0-300ms) for object borders (red) and non-border image elements (blue) when equating the level of V1 activity predicted by VGG-19, the model accounting for most of the variance in V1 activity. Object borders elicit more activity than non-object borders even if the level of V1 activity predicted by the VGG-19 model is equated. The result resembles that obtained when equating RMS contrast (Fig. 4).

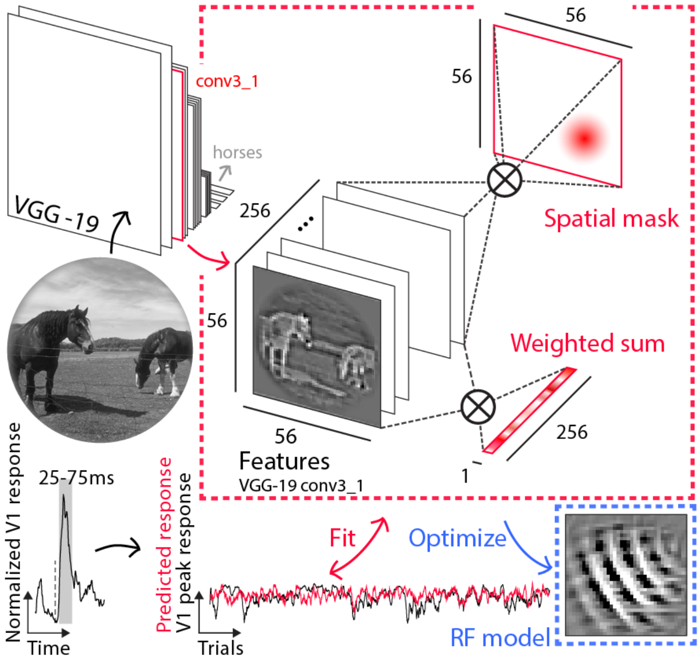


**Figure S9. Schematic of V1 RF model based on VGG-19**

We predicted V1 activity (in a time window from 25-75ms) based on layer conv3_1 of VGG-19. We used the VGG-19 features to predict activity at every recording site with a two-stage mapping (red box; the mapping procedure is described in Methods). The model was inverted to visualize the features to which the neurons were tuned according to this model (blue box).

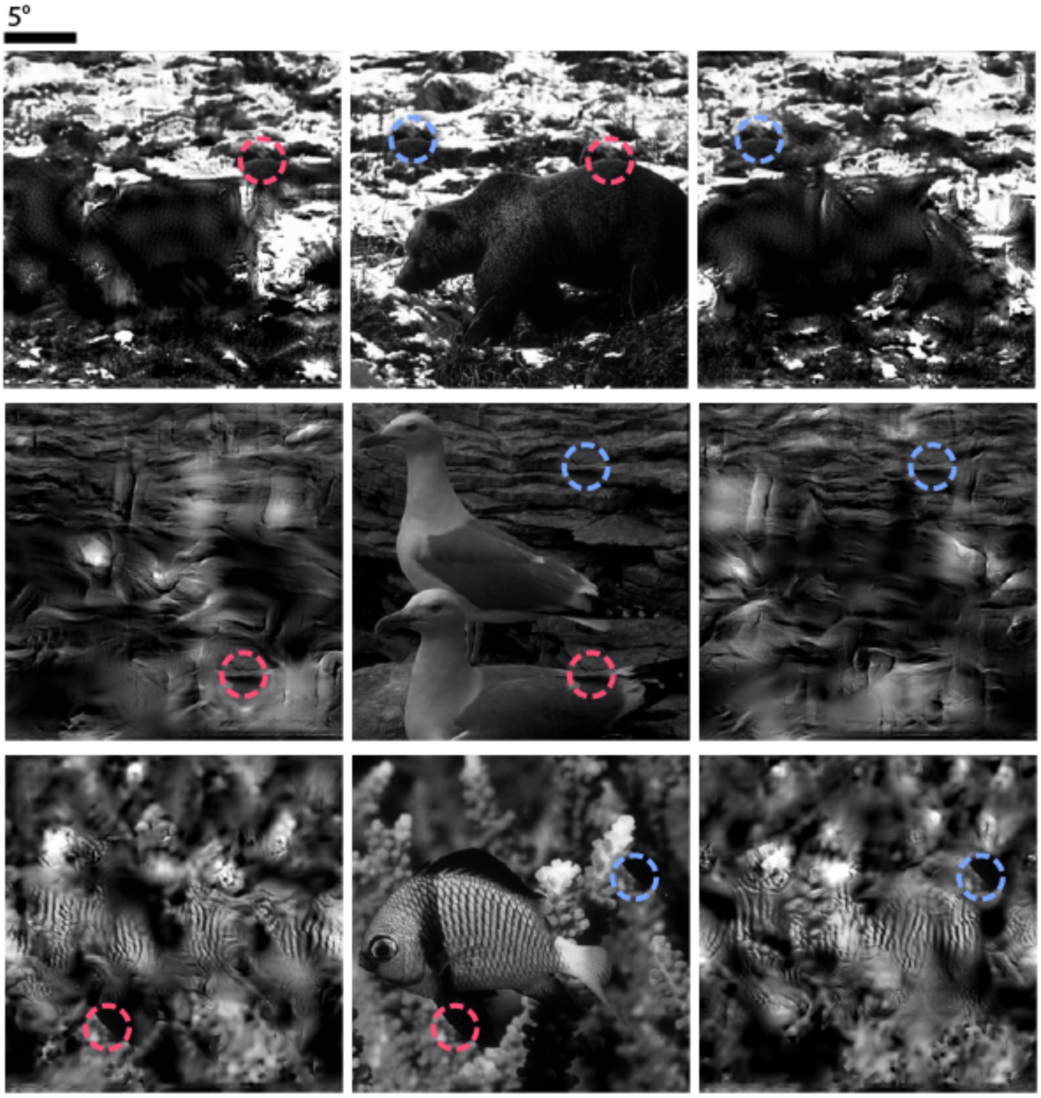


**Figure S10. Original and scrambled stimuli (metamers).**

Example stimuli used in the contextual BoM experiment in which the stimulus in the RF was held constant. Images for which we copied an image patch with an object contour to a background location are shown in the middle. The left and right panels show metamers with a preserved image patch at the object contour (red circles) or the background location (blue circles).
